# Multiomics and digital monitoring during lifestyle changes reveal independent dimensions of human biology and health

**DOI:** 10.1101/2020.11.11.365387

**Authors:** Francesco Marabita, Tojo James, Anu Karhu, Heidi Virtanen, Kaisa Kettunen, Hans Stenlund, Fredrik Boulund, Cecilia Hellström, Maja Neiman, Robert Mills, Teemu Perheentupa, Hannele Laivuori, Pyry Helkkula, Myles Byrne, Ilkka Jokinen, Harri Honko, Antti Kallonen, Miikka Ermes, Heidi Similä, Mikko Lindholm, Elisabeth Widen, Samuli Ripatti, Maritta Perälä-Heape, Lars Engstrand, Peter Nilsson, Thomas Moritz, Timo Miettinen, Riitta Sallinen, Olli Kallioniemi

## Abstract

In order to explore opportunities for personalized and predictive health care, we collected serial clinical measurements, health surveys and multiomics profiles (genomics, proteomics, autoantibodies, metabolomics and gut microbiome) from 96 individuals. The participants underwent data-driven health coaching over a 16-month period with continuous digital monitoring of activity and sleep. Multiomics factor analysis resulted in an unsupervised, data-driven and integrated view of human health, revealing distinct and independent molecular factors linked to obesity, diabetes, liver function, cardiovascular disease, inflammation, immunity, exercise, diet and hormonal effects. The data revealed novel and previously uncovered associations between risk factors, molecular pathways, and quantitative lifestyle parameters. For example, ethinyl estradiol use had a distinct impact on metabolites, proteins and physiology. Multidimensional molecular and digital health signatures uncovered biological variability between people and quantitative effects of lifestyle changes, hence illustrating the value of the combined use of molecular and digital monitoring of human health.

---

Longitudinal measurements of clinical laboratory tests, multiomics profiles, and digital health monitoring using wearable sensors could facilitate a comprehensive analysis of human health and the impact of lifestyle and disease. This provides opportunities to explore human systems biology and to predict and intervene in processes leading to disease. However, our ability to make use of such data and define actionable insights, correlations and causal patterns is limited. There is a shortage of deep integrated datasets, where different dimensions of human biology have been explored at the same time and their changes over time recorded and/or connected with digital monitoring data from wearable devices. Each individual has a unique profile, composed of genetic, epigenetic, molecular, clinical and lifestyle parameters, which may change over time during development, aging and disease transitions. These processes cannot be understood as a single general pathway for the average human or patient, but they can be better described as a collection of individual health trajectories, where changes in individual variables are highly interconnected. Several studies have investigated the use of individualized molecular profiles to assess disease risks or the connection between omics measurements and clinical tests, using an n-of-one longitudinal approach^1,2^, a cross-sectional design^3^, a controlled longitudinal perturbation study^4^, and a cohort study with personal behavioural coaching^5^. More recently, other studies^6–8^ have analysed longitudinal data over an extended time period in a cohort of individuals at risk for diabetes, while another prospective observational study investigated the stability of the individual molecular signatures^9^.

The Digital Health Revolution (DHR) program^10^ was based on the concept that future healthcare strategies will evolve in a direction which allows citizens to control and make use of their personal data to improve their health and wellness. The project aimed at implementing proactive P4 healthcare (predictive, preventive, personalized, and participatory), with multi-level molecular and digital data. Within this framework, we integrated deep multiomics profiles and connected them to health surveys, clinical observations and digital health measurements. We aimed to 1) integrate longitudinal multi-omics and digital health data between and within people over time to reveal subgroups and personal trajectories; 2) exploit the discovered associations to understand novel links between molecular, clinical and digital health data and 3) verify if data feedback and coaching would guide and motivate people to undertake lifestyle changes. Overall, we aimed at achieving a holistic understanding of the parameters involved in different aspects of human biology and health. To accomplish these goals, we applied multiomics factor analysis and then connected the learned factors with interpretable characteristics of health and behaviour as well as quantitative digital health data.

## Results

### Study overview

We carried out a 16-month longitudinal study on 96 individuals (aged 25-59) recruited from an occupational healthcare clinic in Helsinki, Finland (Supplementary Figure 1-2 and Supplementary Table 1). The participants had no previously diagnosed serious chronic diseases, although we allowed individuals with risk factors for chronic diseases to participate. We acquired comprehensive measurements of health and behaviour, including anthropometrics, clinical laboratory tests, questionnaire data, physical fitness tests, and activity and sleep quantification with a wearable device. In addition, we performed a series of omics measurements to study the genome, the plasma proteome and metabolome, the autoantibody profile and the gut microbiome. The prospective collection of molecular and digital profiles resulted in a thorough longitudinal dataset of human health and lifestyle (Figure 1). We defined baseline molecular profiles and longitudinal data trajectories in the participants during personalized lifestyle coaching. We collected over 20,000 biological samples during five study visits and generated a compendium of >53 million primary data points for 558,032 distinct features. Two types of feedback were applied to stimulate and motivate lifestyle changes. First, actionable health data were returned to participants via a web dashboard and interpreted by a study physician, starting from the second visit. Second, personal data-driven coaching was provided, including three face-to-face and six remote meetings, plus continuous email and phone support, starting from the third visit. Personal actionable possibilities were identified with the help of questionnaires, clinical laboratory tests and physical examination and focused on diet, exercise, mental wellbeing, stress and time management. Out of the 107 people enrolled, 96 completed the study. The data-driven coaching positively impacted on the health of the participants, and 86% reported improvement in at least of the following aspects: diet, exercise, sleep, mental wellbeing, stress management, drinking or smoking (Supplementary Figure 3). For example, the percentage of people who did not exercise decreased from 9% to 1%, the percentage of smokers decreased from 16% to 8%, and the percentage of daily drinkers from 8% to 5%. A major achievement was the decrease in the prevalence of vitamin D deficiency (31% to 16%, Sallinen et al. The Journal of Nutrition, in press). The participants described the wearable device, the return of personal health data and the tailored coaching as the most motivating aspects of the study (Supplementary Figure 3). Overall, the subjects were enthusiastic about participating and 76% indicated that they would again participate in a similar study.

**Figure 1:**
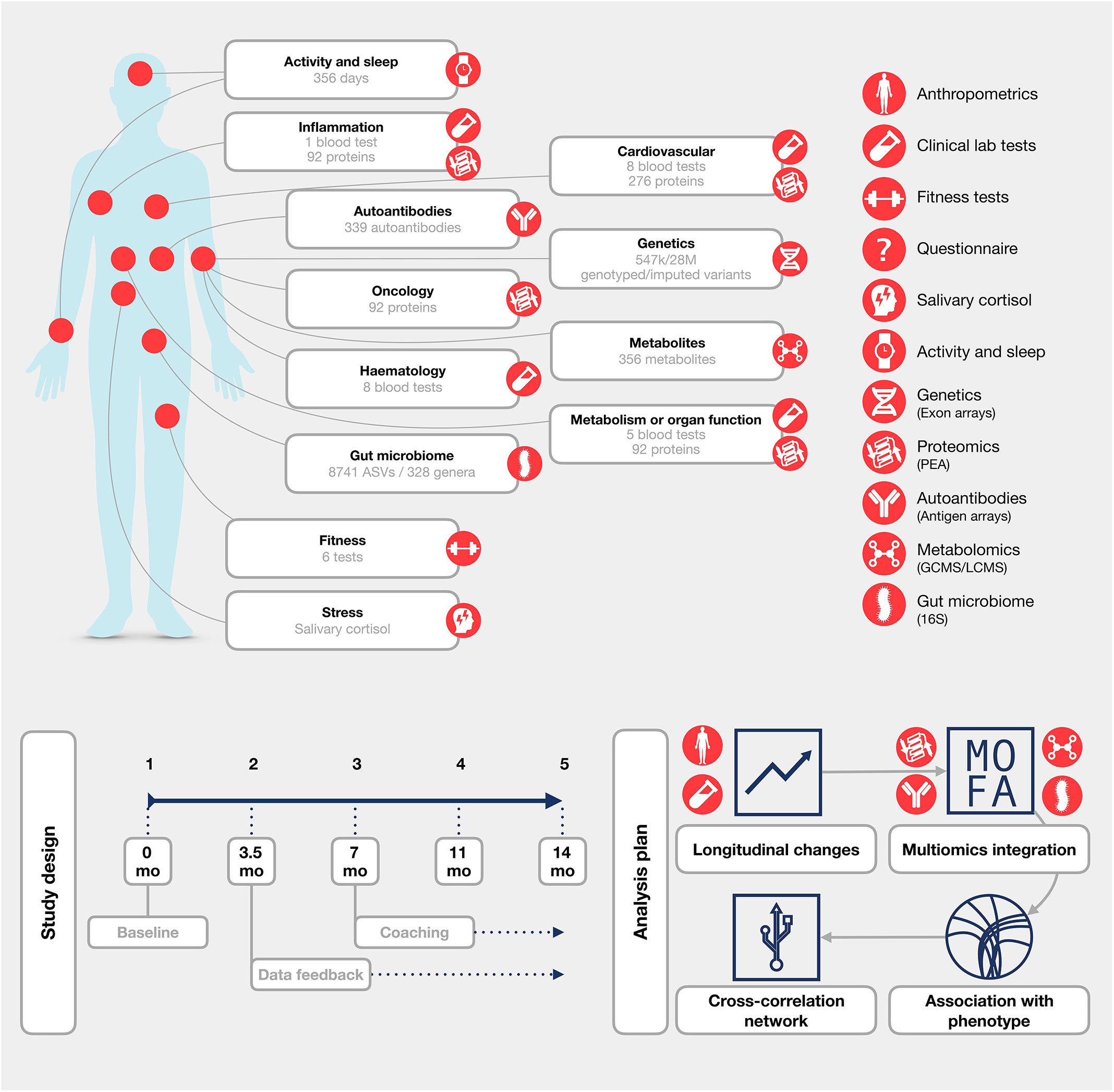
Overview of the measured features, longitudinal design and analysis workflow.

### Longitudinal actionable changes

We defined a *phenotype* using the ensemble of questionnaires, anthropometrics, vitals and clinical laboratory tests. Actionable health issues were defined at baseline, including dyslipidaemia, overweight and obesity, elevated systolic blood pressure, low vitamin D, elevated blood glucose, anaemia and self-reported obstructive sleep apnea (Supplementary table 1 and Supplementary Figure 4).

We first analysed the clinical measurements at baseline to define the group of individuals with Out-Of-Range at Baseline (OORB) values. To explore if data feedback and coaching had a positive effect on objective health parameters, we modelled the longitudinal changes in the ORRB individuals, based on the assumption that measurements outside the reference values represented an actionable finding to improve lifestyle and health. We analysed the average change per visit, adjusting for sex and age. We found significant improvements (FDR<0.05) in several key health parameters for the OORB group during the course of the study, such as an increase in vitamin D levels and a decrease in blood pressure, LDL-cholesterol, and total/HDL cholesterol ratio (Figure 2 and Supplementary Table 3).

**Figure 2:**
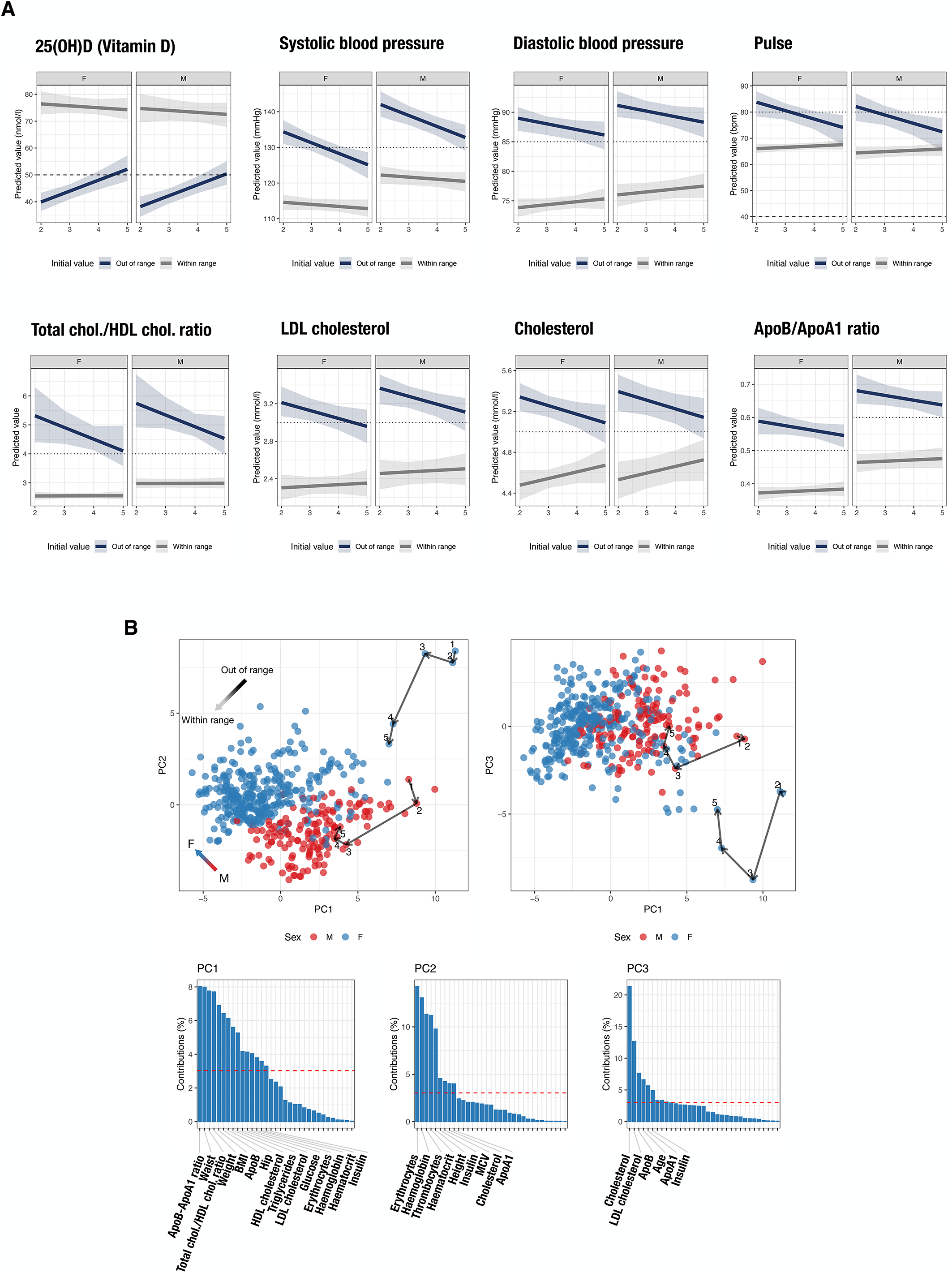
**A.** Longitudinal analysis reveals the positive effect of data feedback and coaching on health. Generalized Estimating Equation (GEE) model prediction for the significantly changed variables (FDR<0.05), stratified by sex and out/within range status at baseline. For this analysis, the second visit represented a baseline because data feedback and coaching started at the 2nd and 3rd visits respectively (grey: within range, blue: out of range). Bootstrapped confidence intervals are shown. **B.** Principal components plot of the individuals obtained using all the numerical clinical variables. The trajectories for two selected individuals along successive study visits are shown with arrows. The loadings for the first three PCs are shown in the bottom panels and the variables with relative contribution higher than average (red line) are named.

Secondly, we obtained an overview of all the clinical variables and compared the individuals using Principal Component Analysis (PCA). The first three PCs (Figure 2) accounted for 44.8% of the variance and the major drivers of variability were cardiometabolic variables (for PC1 and PC3: BMI, insulin, fasting glucose, cholesterol, and lipoprotein profiles) and sex-dependent anthropometrics and haematological measurements (PC2). Trajectories for selected individuals showed the changes during subequent study visits and visualized the improvements in the clinical parameters (Figure 2 and Supplementary Figure 5).

In summary, we showed that the return of data as well as data-driven lifestyle coaching resulted in objective improvements of health behaviour and health outcomes, including physiological and laboratory measurements indicative of cardiovascular risk.

### Multiomics signatures

#### Distinct sources of molecular variability

We next generated multiomics profiles and set out to explore the association of these variables with each other and with the clinical features. We first investigated the sources of variability in the omics, by partitioning the variance for each feature into personal variation (i.e. between-individual), known factors (age, sex and study visit) and residual unknown sources (for example biological, environmental or technical aspects). We found that the personal variation impacted all data types but to a different extent (Supplementary Figure 6). Autoantibodies represented a highly personal signature and on average 92% of the variance was explained by personal variation. Metabolites were dominated by personal variation (56-71% on average) but residual components were also present (23-35% on average). Proteins had similar average personal (49%) and residual components (45%). The microbiome data was dominated by unexplained factors (68% on average). These results were confirmed by an alternative distance-based method (Supplementary Figure 6). We observed that the average within-individual distance was lower than the between-individual distance, to a different extent for different omics, and autoantibodies showed the lowest within-individual distance, i.e. high similarity.

### Understanding the molecular variability

We used Multi-Omics Factor Analysis (MOFA+)^11,12^ to carry out an unsupervised analysis of the complete multiomics data, hence excluding the clinical variables and other phenotypic data. MOFA+ helped to integrate and interpret the measurements and their variability across all the omics layers. The analysis resulted in an interpretable low-dimensional representation in the form of a small number of independent factors that captured the major sources of variability in the data and originated from simultaneous inclusion of baseline and longitudinal differences. The analysis facilitated the identification of subgroups of samples and the molecular features that contributed to the ordination of the samples in the dimensions defined by the factors. An overview of the samples based on the learned factors is given in Supplementary Figure 7. We identified 14 factors (F1-F14), each explaining a minimum of 2% of the variance in at least one data type. The fraction of the total variance explained across the six omics types varied from 6% for gut microbiome to 95% for the autoantibodies (Figure 3). Some of the learned factors were predominantly associated with one omics type, while others contributed to several omics, suggesting that the underlying biological determinants might affect different types of measurements simultaneously. We then explored the molecular basis of each factor. First, we investigated the contribution of the original omics features, by inspecting the loadings on each factor, which represent their weights, in order to interpret the nature of the associated biological domains. Second, we tested the association between the factors and the phenotype, including questionnaire data, clinical variables, fitness tests, activity and sleep data, and other metrics. Each factor was linked to specific phenotypic characteristics (linear models, FDR<0.001, Figure 3 and Supplementary Table 4). Importantly, most of the factors were associated with distinct sets of variables across the different categories and thereby defined molecular patterns that were linked to distinct aspects of lifestyle and health, including modifiable behavioural aspects, such as diet and exercise. We then refined the top associations for each factor with a covariate-adjusted statistical model. We aimed at accounting for the correlation among observations from the same individual and fitted a linear mixed model (LMM), with age and sex as fixed effects and the individual as a random effect. We also interpreted the association in view of the contributing molecular features and pathways and the results are presented below.

**Figure 3.**
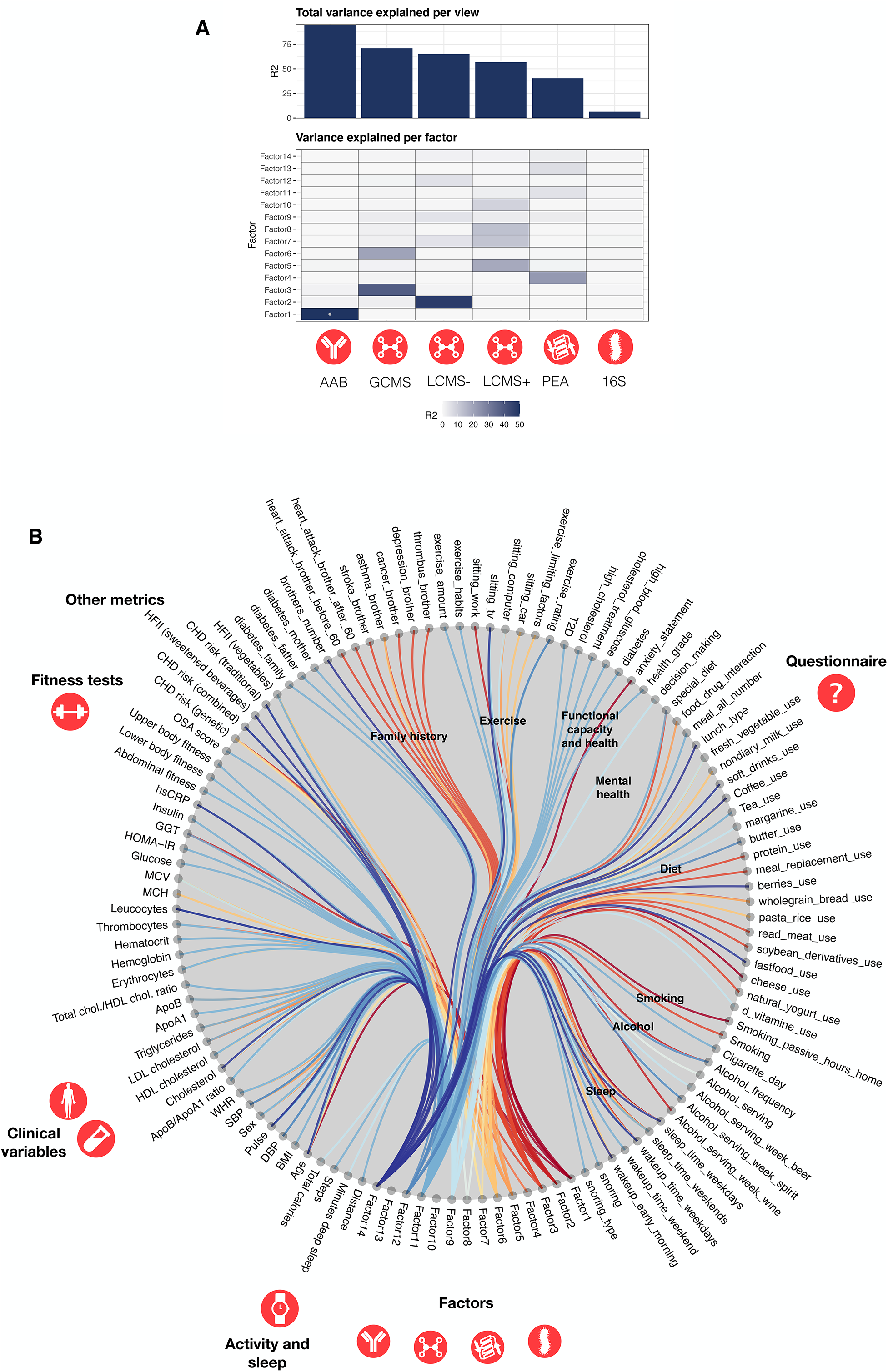
**A.** Variance decomposition. Percentage of total variance explained (R^2^) for each data layer (up) or variance explained (R^2^) for each factor and data layer (bottom). A dot marks the out-of-bound value (>50%) for autoantibodies. **B.** Association with phenotype. The factor values were tested for their association with the phenotypic variables and the top associations (FDR<0.001) are shown. The edges are bundled according to their category and colored based on the originating factor.

#### Obesity and insulin resistance

The top features with high positive loadings on F11 included plasma proteins involved in pathways known to be dysregulated in obesity or dyslipidemia (LEP, IL-1ra (IL1RN), t-PA (PLAT), FABP4, FGF21, IL-6, LDL receptor)^13–18^. Interestingly, several proteins reported as protective factors against obesity (GH1, IGFBP1, PON3)^19–21^ had negative loadings. We found significant correlations between F11 and well-known metabolic traits and clinical measurements (Figure 4 and Supplementary figure 9). These included positive association with BMI (p_lmm_=1.53E−21), LDL-cholesterol levels (p_lmm_=4.47E−03) and insulin resistance (log10HOMA-IR, p_lmm_=6.51E−14). Also, we found a negative association with HDL-cholesterol (p_lmm_ =2.16E-06) as well as with measures of physical activity and fitness gathered from the questionnaire, such as leisure-time exercise (p_lmm_=1.06E−03) and exercise habits (p_lmm_=8.25E−04), or the activity levels in terms of steps/day measured by the wearable device (p_lmm_=0.01). An inverse correlation was also observed with physical fitness at the level of upper body (p_lmm_=5.53E−05) abdominal (p_lmm_=1.59E−03) and lower body (p_lmm_=1.14E−03). Participants with first-degree relatives with diabetes had higher factor values compared to the individuals who did not report family-history for diabetes (p_lmm_=6.25E−03). Finally, an individual who was diagnosed with type 2 diabetes during the study had the highest factor values. We therefore interpreted F11 as strongly related to obesity, insulin resistance and diabetes risk and pathogenesis as well as clearly associated with quantitative data on exercise and mobility. F11 was also associated with overall health, as evidenced by a negative association with self-reported overall health (p_lmm_=9.26E−07) and a positive correlation with coronary heart disease (CHD) risk score (p_lmm_=0.01). In summary, F11 assisted in estimating molecular patterns of health and behaviour that are associated with obesity as well as potential trajectories leading to diabetes.

**Figure 4.**
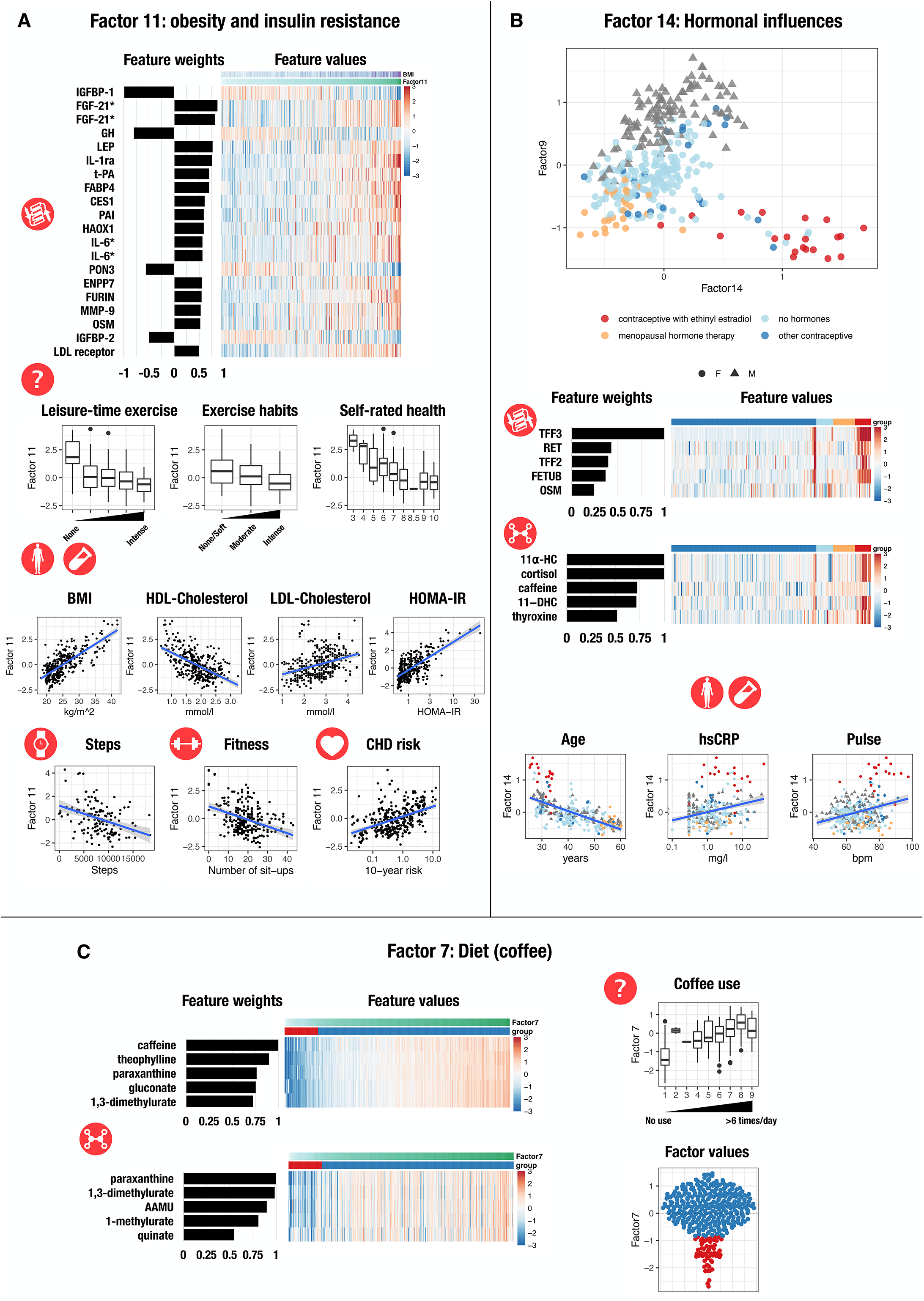
**A.** Factor 11 is linked to obesity and insulin sensitivity. The top scaled protein weights for Factor 11 are shown as well as their values (row z-score). The association between the factor values for each sample and the phenotypic variables is shown. A regression line with 95% confidence interval is shown when appropriate. An asterisk marks the proteins measured on multiple PEA panels. **B.** Factor 14 is influenced by hormones. A scatterplot of Factor 9 and Factor 14 values shows that samples can be separated based on the sex (Factor 9) and the use of hormones (Factor 14). Factor 14 identifies a group of young women using contraceptives with ethinyl estradiol (red). The top metabolites and proteins with positive scaled weights on Factor 14 are shown. The original feature values are shown as a heatmap (row z-score). The association between the factor values for each sample and the phenotypic variables is shown, as well as a regression line with 95% confidence interval. Points are coloured as in the top panel. Abbreviations: 11⍺-HC: 11alpha−hydrocortisone, 11−DHC: 11−dehydrocorticosterone. **C.** Factor 7 is associated to coffee consumption. The scaled positive weights are shown for the top metabolites, as well as their values. The association between the factor values with self-reported coffee consumption is shown. A two-group classification of the samples based on Factor 7 values was obtained with gaussian-mixture modeling and shown with colored dots (red: lower coffee consumption, blue: higher coffee consumption).

#### Impact of hormones and ethinyl estradiol

F9 was influenced by sex and sex-specific biological differences in multiomics. In contrast, F14 was linked to the use hormone replacement therapy or hormonal contraception in women. Young women reporting the use of a contraceptive containing ethinyl estradiol (EE) had particularly high F14 values (p_lmm_=2.28e−10). Consistent with this observation, known estrogen-sensitive plasma proteins had the highest positive weights for F14 (Figure 4), but there were also contributions from cortisol and thyroxine levels, which we interpreted as potential secondary effects^22,23^. Pulse (p_lmm_=9.32E−03), hsCRP (log-transformed, p_lmm_=7.40E−09) and leukocyte numbers (log-transformed, p_lmm_=6.09e−05) also positively correlated with F14 and alterations of these clinical variables were prominent in women with particularly high factor values. These observations indicated, in an unsupervised manner, that EE use is associated with strong molecular and physiological effects on human biology, including increased levels of inflammation, impact on thyroid, cortisol levels and pulse. These effects are distinct from all the other sources of variability in our study population,

#### Dietary habits

F3 and F6 were mostly influenced by metabolites and associated with distinct dietary habits assessed with the questionnaire (Supplementary figure 10), such as consumption of fresh fruit and vegetables. However, when accounting for the effect of age, sex and individual, the dietary characteristics were not significantly associated with the factors (p_lmm_>0.05), suggesting that the associations may be explained by individual-specific (person-to-person) differences in dietary patterns and not by changes in the diet during the study. Moreover, F7 was largely linked to caffeine use as determined both by the molecular nature of the features and the associations with questionnaire data. Indeed, caffeine and its metabolites had the largest loadings for F7 (Figure 4), and this factor was strongly correlated with self-reported coffee consumption (p_lmm_=7.63E−12).

#### Lipids, free fatty acids and fatty acid esters

F5, F8 and F12 (Supplementary figure 11) were each reflecting distinct aspects of the plasma lipid composition. Free fatty acids (FFA) contributed mostly to F12, while fatty acid esters (FAEs), mostly in the form of acylcarnitines, contributed to F5 and F8. In the postabsorptive state, blood FFA are a result of lipolysis from adipose tissue and they reflect dietary intake^24^, while changes in the acylcarnitine pool can be linked to changes in fatty acid oxidation. Accumulation of acylcarnitines as products of incomplete mitochondrial fatty acid oxidation has been associated with obesity and diabetes^25^. Even after adjusting for the covariates, F12 was negatively associated with blood pressure (systolic, p=0.035; diastolic p=0.021), in line with the known association between FFA and hypertension^26^. F5 and F8 were both strongly associated with clinical measurements indicating dyslipidemia, including total cholesterol (p_lmm_=1.64E-08 and p_lmm_=8.48E−10) LDL cholesterol (p_lmm_=7.13E-06 and p_lmm_=1.74E−05), triglycerides (p_lmm_=3.74E-05 and p_lmm_=8.45E−7) and ApoB (p_lmm_=3.24E-08 and p_lmm_=9.11E−06), thus implicating dysregulation of lipid metabolism as an underlying influence for F5 and F8 Importantly, F5 and F8 were only weakly associated with BMI (R2=0.004 and R2=0.006; p_lmm_=0.05 and p_lmm_=0.07), suggesting that these two factors were associated with dyslipidemia independently of obesity.. Sex was a significant covariate for the associations seen for F5 (maximum p_lmm_=1.21E-05) but not for F8 (minimum p_lmm_=0.73).

#### Hepatic function

F2 was influenced by metabolites connected to liver function (Supplementary figure 12). Bile acids or their glycine or taurine conjugates had negative weights and a major impact in the classification of the individuals along the F2 axis. Furthermore, F2 included liver-associated metabolites that on average were increased in the samples with negative factor values. These metabolites are known to be increased after liver disease (biliverdin) or are known to be used to eliminate hepatic nitrogen pool (glutamine, hippuric acid, phenylacetylglutamine). The liver is a fundamental organ in maintaining metabolic homeostasis, it modulates the composition of the blood metabolome and therefore we assumed that F2 reflected the structural and functional integrity of the liver.

#### Autoantibodies

F1 was mostly explained by the presence of autoantibodies, it was influenced by the binary nature of the autoantibody data (presence/absence) and by the stable autoantibody signature over time. Indeed, autoantibodies can be considered as an IgG reactivity barcode for each individual^27^. F1 correlated well with the autoantibody counts and to a lesser extent with the age (Supplementary Figure 8), in agreement with the fact that autoreactivity is more common in the elderly, possibly linked to age-associated B cells^28^.

In summary, we demonstrated that we could dissect the overall pattern of molecular variation between and within individuals using the multi-omics factor analysis. Each of the 14 factors reflected distinct aspects of human biology, behaviour, lifestyle, use of drugs (hormones) or potential transitions to disease.

### Correlation networks: linking molecular features with clinical variables, questionnaire data and digital health monitoring

We then performed correlation analysis, including all the multiomics data as well as an ensemble of quantitative or semi-quantitative measures consisting of clinical data, activity and sleep, fitness tests, and other scores. We computed a cross-correlation network between features of different types using two alternative metrics. For the between-individuals network (BIN), we averaged the measurements from all time points before calculating correlations. We also included 146 genetic trait scores obtained by summing up the contribution from all the variants associated with each trait as reported by the GWAS catalog. The within-individuals network (WIN) consisted of linear correlations of repeated measures of pairs of features within-participants. It estimated a common regression slope, representing a measure of association shared among individuals, and resulting from changes occurring at the individual level during the study. The BIN and WIN were calculated after age- and sex-adjustment and p-values were corrected for multiple hypothesis (FDR<0.05). We then identified distinct communities of interconnected features (modules), as an aid for interpretation.

Among the strongest correlations of the BIN (|ρ| > 0.6), we detected distinct measurements of the same molecular entity using different assays (i.e. TSH (clinical-PEA), ρ= 0.96; Cholesterol (clinical-GCMS) ρ=0.82), or associations between different features such as LDL-receptor protein and triglycerides (ρ=0.82), or LEP protein and BMI (ρ=0.68). The network consisted of 375 nodes and 570 edges (Supplementary table 5). Several clinical variables represented hubs with the largest number of associated connections to metabolites and proteins (Figure 5). These hubs were often inversely correlated with modifiable behavioural risk factors, such as wearable-based measures of physical activity (e.g. insulin with intense physical activities, waist circumference with number of steps). Similarly, measures of physical fitness (lower body, abdominal and upper body) correlated inversely with plasma LEP levels. Notably, the proteins with common genetic defects in familial hypercholesterolemia appeared in the cardiometabolic subnetwork (LDLR, PCSK9, and ApoB). For instance, PCSK9, a pharmacological target of LDL-lowering therapies, correlated positively to several glycerophospholipids and ApoB, but negatively to CMPF (3-Carboxy-4-methyl-5-propyl-2-furanpropionic acid), a metabolite that is formed from the consumption of fish oil and may have positive metabolic effects^29^. The whole network included 24 GWAS summary scores (Supplementary figure 13), and 11 scores concerned haematological measurements (erythrocytes, leukocytes and thrombocytes). We observed a direct link between the genetic score and the corresponding trait measured in this study, including haematological traits, but also associations between proteins or metabolites and the genetic susceptibility to a disease/trait with cardiometabolic relevance. For example, we observed an inverse association between the genetic risk score for BMI in physically inactive individuals and several FAEs, carnitine and tyrosine. Furthermore, an inverse association was observed between the genetic risk score for abdominal aortic aneurysm and IL-6RA levels, supporting the contribution of IL-6 signalling and inflammation in the pathogenesis of this disease^30^. These observations indicate that the between-individuals variability may be explained at least partly by personal genetic differences and that these genetic differences exert their impact via specific molecular pathways.

**Figure 5.**
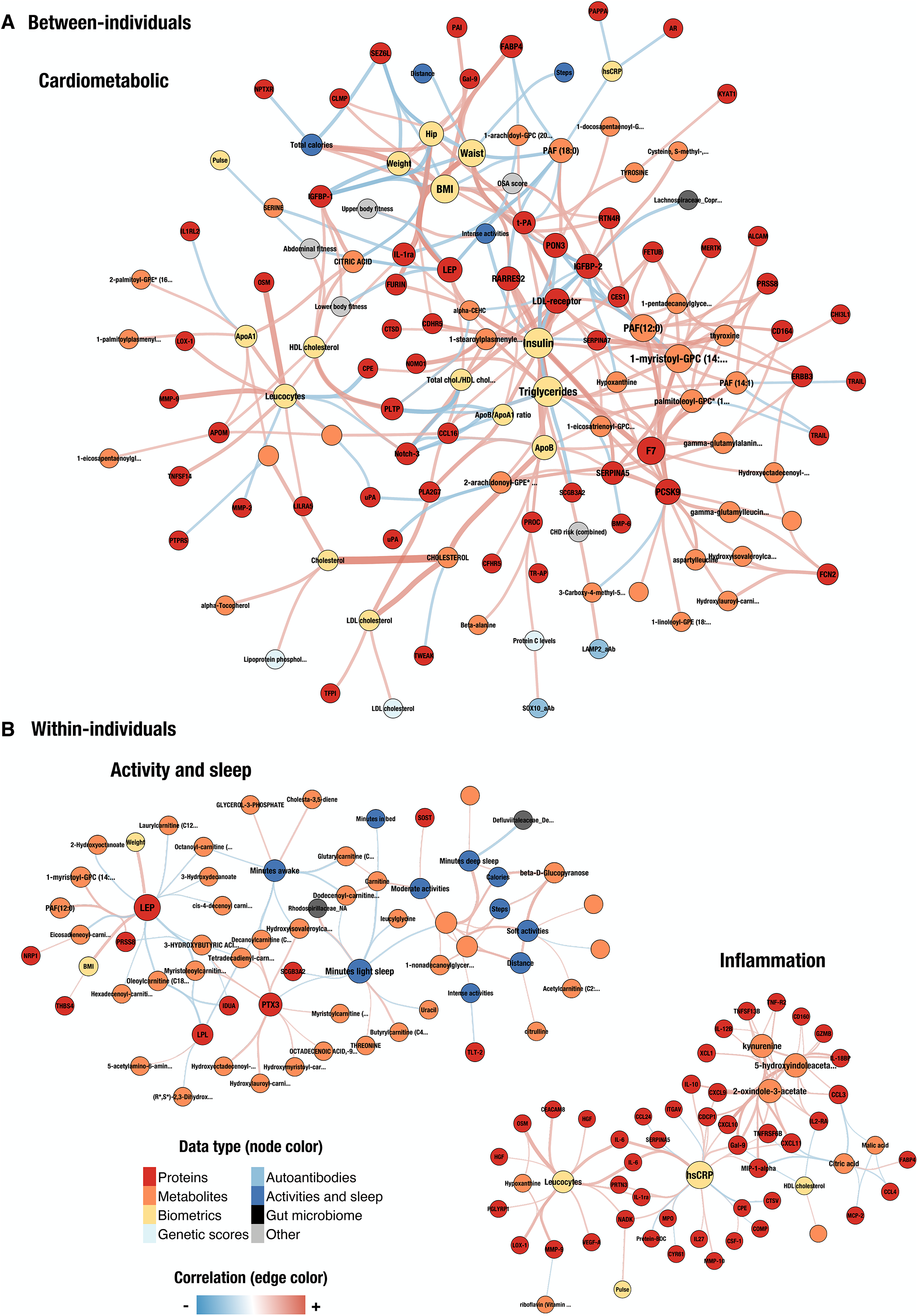
Cross-correlation network analysis aids the interpretation the relationship between data types. Only associations between features of different type were considered to calculate correlation networks at FDR<0.05. The size of the nodes is proportional to the node degree and the edge thickness is proportional to the magnitude of the correlation. The node color indicate different data types. The proteins appearing twice in the network were measured on multiple PEA panels. **A.** Selected between-individual subnetwork. **B**. Selected within-individual network modules.

The WIN captured associations between pairs of features that resulted from concordant variation in repeated measures obtained from the same individuals. This source of variation is assumed to be complementary to the BIN, it does not average or aggregate the data, neither does it violate the assumption of independence of the observations. Thus, the edges in this network resulted from changes occurring over time and not due to baseline between-individuals characteristics. The network consisted of 302 nodes and 611 edges (Supplementary table 5). Only 46 edges and 151 nodes were also detected in the BIN, reflecting how BIN and WIN highlight non-overlapping correlations, i.e. they reveal distinct types of biological associations. For example, in an inflammation-related subnetwork (Figure 5), hsCRP and leukocytes lay in the proximity of cytokines, cytokine receptors, proteins involved in leukocyte functions and members of the tryptophan metabolic pathway (kyneurine and indoles). This subnetwork was not revealed in the BIN and the observed edges likely originate from individuals transitioning through inflammatory states or infections during the study, although other sources of variations (season, exercise etc.) cannot be excluded. Other edges specifically observed in the WIN showed metrics of activities and sleep in the proximity of several metabolites and proteins. Among the proteins, we observed well-known members of lipid metabolism pathways (LEP, LPL) and proteins with related functions (PTX3^31,32^, NRP1^33^, PRSS8^34^). Correspondingly, FFA and FAEs appeared in this subnetwork. These correlations suggest that changes in dietary factors, fat metabolism, physical activity and sleep behavior as a result of the lifestyle coaching could be further investigated as possible explanations for the observed effects.

### Personal trajectories

In order to stimulate lifestyle changes and improve health, we defined personalized, actionable data-driven possibilities for lifestyle guidance for each participant. This strategy resulted in subgroups where similar changes in actionable clinical variables were present. When considered at the level of individuals, the changes unfolded into personal trajectories, and often a connection with the patterns in multiomics data were established. Changes in diet and exercise were common among participants, as illustrated by the case of a 28-year male, where the tailored lifestyle change (nutrition and exercise) resulted in an improvement of clinical variables and self-rated health (Figure 2 and Figure 6). At baseline, he presented with obesity, high systolic blood pressure, dyslipidaemia, elevated insulin levels and elevated hsCRP (Figure 6A). During the study, he reported increases in self-rated health, frequency and amount of exercise (Figure 6C). The latter was also objectively confirmed by the activity levels detected by the wearable device (Figure 6B). The sustained level of activity resulted in longitudinal changes in key actionable clinical variables at the end of the study (Figure 6A), including decreased BMI, improved lipid profile, reduced low-grade systemic inflammation, reduced systolic blood pressure, normalization of insulin levels and improvement of the estimated insulin resistance. These interpretable changes reflected in molecular changes, as shown by the MOFA+ factors (Figure 6D). Indeed, the improvement in BMI, lipid profile and insulin resistance, combined with an increase in physical activity, was reflected as a decrease in F11 values. We observed decrease of obesity- and inflammation-associated proteins and other adaptations suggesting an involvement of the GH/IGF-1 axis. Similarly, the changes in F8 and F10 - associated with fatty acid esters and lipid metabolism - reflected the variation in the total cholesterol levels. Several acylcarnitines peaked at the third visit, suggesting a metabolic adjustment during a transition period toward increased physical activity. For comparison, we show (Figure 2 and Supplementary figure 14) the case of a 36-year-old female, presenting with obesity, dyslipidaemia, hyperinsulinemia and hyperglycaemia at baseline. The study protocol required a referral to a physician, and she was indeed diagnosed with type 2 diabetes. During the study, phenotypic changes occurred: reduction in the glucose, insulin and cholesterol levels, as a result of a new therapeutic intervention (Metformin, Atorvastatin, Empagliflozin), while she did not undertake a genuine lifestyle change and did not increase physical activity. Interestingly, the F11 levels did not change much, this individual remained an outlier at all the time points and we did not observe a trend in the corresponding molecular features indicating reduction of inflammation and adiposity.

**Figure 6.**
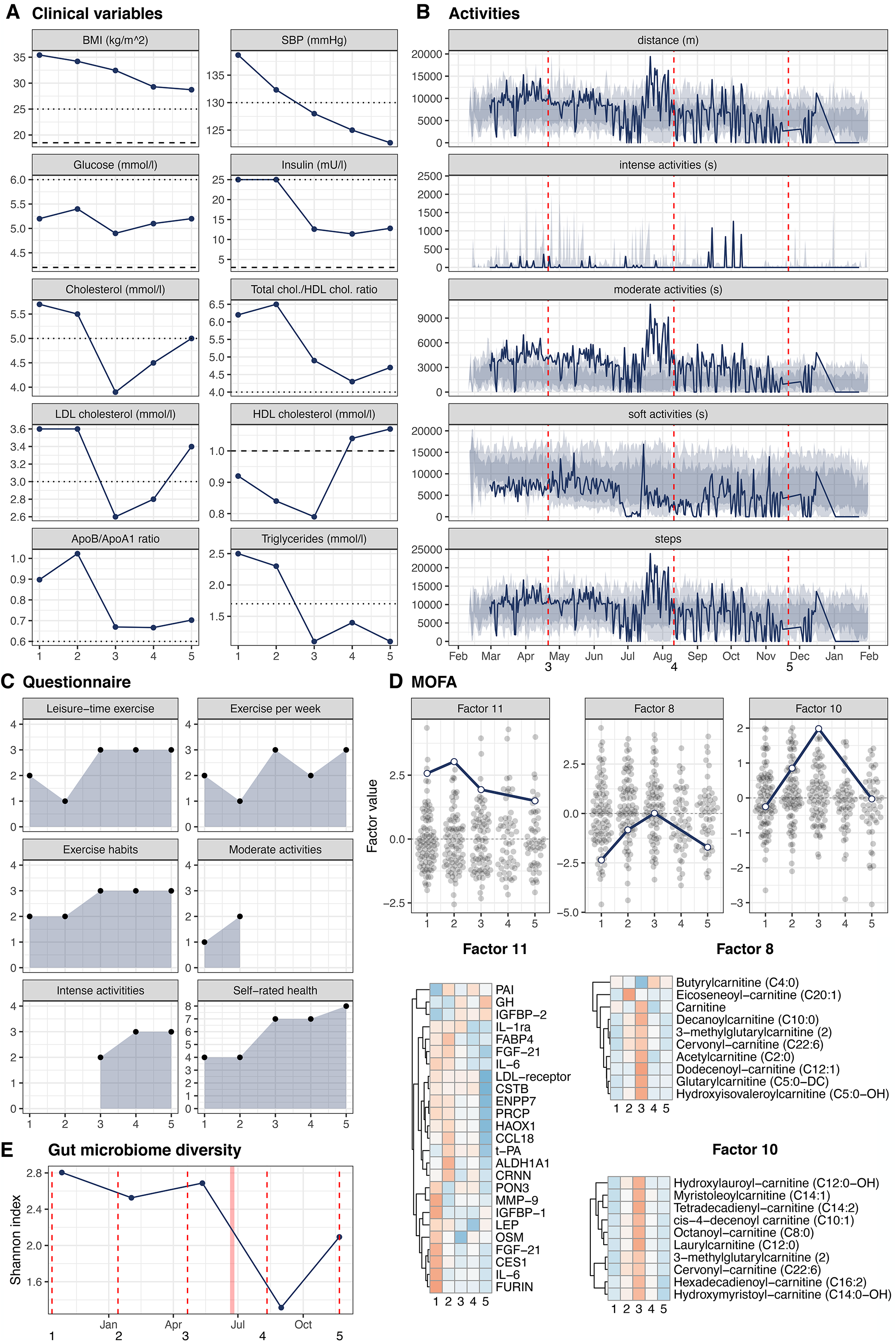
A case study: lifestyle change in one individual and the associated molecular changes. **A.** Clinical variables showing longitudinal changes. Reference values are shown (dashed line: lower normal value, dotted line: upper normal value) **B.** Daily summary of the activities recorder by the smart watch included: distance, time spent in intense, moderate or soft activities and number of steps. The shaded area correspond to the 10, 25, 75 and 90 percentiles in the whole population. The study visits are marked with vertical dashed lines **C.** Selected lifestyle questions. The numbers correspond to orderer categories, where a higher number corresponds to higher frequency or more favorable outcome. **D.** Longitudinal values for selected MOFA+ factors (blue line) plotted together with the factor values for the remaining samples at each study visit (grey dots). The top features for the corresponding factors are shown as heatmaps (row z-score, blue:low, red:high). **E.** Gut microbiome diversity (Shannon index). The vertical dashed line mark the date for the study visit, which could be different than the fecal sample collection date (dots). A red shaded area mark a period of antibiotic treatment.

A case of a participant with a respiratory tract infection is illustrated in Supplementary Figure 15, it shows the dysregulation of metabolic pathways during infection as previously observed^1,6^, and highlights how metabolic signatures can be prominently altered during infections.

## Discussion

The convergence of systems medicine, digital health, big data, consumer-driven healthcare and social networks is at the heart of the P4 healthcare. To facilitate the adaptation of the P4 approach into the current healthcare systems, longitudinal health intervention studies are needed. Indeed, numerous initiatives exist to profile healthy and diseased individuals with clinical features, multiomics and digital health monitoring^1–8^. However, fewer studies have simultaneously combined longitudinal multiomics profiles, digital health data monitoring, data feedback and tailored lifestyle coaching. Here, we report a truly comprehensive approach to generate longitudinal data from 96 people who underwent tailored lifestyle coaching for over 16 months. Our study provides several steps to promote an understanding of human biology and health over time and, at the same time, a data-driven, individualized approach to motivate and monitor the impact of lifestyle changes. Firstly, our data emphasize the importance of multiomics profiles in preventive and personalized medicine approaches. Most of the published large-scale medical research initiatives have explored predominantly the impact of genetics on health parameters. We think that the precision medicine toolbox should include multiomics and digital health readouts, longitudinal measurements for each individual. Secondly, we have shown that by integrating all the data in an unsupervised analysis we can rediscover known associations between molecular features and lifestyle changes, health behaviours, adiposity, diet, physical activity or hormonal use as well as identify numerous novel associations. We have also shown how data from a wearable device correlate with changes in specific molecular pathways, which could help to understand the molecular effects of e.g. exercise and sleep on specific aspects of human biology. Thirdly, our data-driven feedback and individualized lifestyle coaching stimulated behavioural changes and we anticipate that this model could become broadly applicable in the future. In this study, genetics, questionnaire and clinical measurements were returned to the participants in real-time, while the full multiomics data were analyzed retrospectively. The analysis of all molecular data at the end of the study also helped to ensure lower cost, higher quality and consistency, and elimination of batch effects.

Our data analysis resulted in a low-dimensional space to explore the complexity of the multiomics and the connection to known sources of variation. Among those, sex, age and BMI are among the most studied and often are confounders in association studies. For this reason, we regressed out the effect of age and sex in the correlation networks, but we allowed the effect of these covariates to appear in the MOFA+ analysis, in order to verify if separate, independent axes of variability could be retrieved. Indeed, we verified that BMI, as a proxy for adiposity and obesity, could only be strongly associated with one factor (F11) and that there was no strong significant association of obesity with any of the other factors. Similarly, sex was associated only with a few factors (F5, F9, F11). Therefore, the identified axes of variation represented a spectrum of molecular variation that could be attributed to a variety of lifestyle factors and could not simply be explained by the confounding effects of obesity, age or sex. This strongly supports the power of our analysis to extract independent axes of molecular variability of potential etiological and pathogenetic importance, and hence with opportunities for unbiased health monitoring.

An example of a highly interesting and novel observation is that the use of oral contraceptives containing EE in a subset of the female participants was alone sufficient to influence the covariance of the proteome and metabolome. The effect was so distinct that F14, as uncovered from this unsupervised MOFA analysis, was almost completely due to the use of this one hormonal drug. We suggest that the strength of this association could have been even greater if we had detailed information on the days of the administration of the hormone and the days of the menstrual cycle of all females when samples were obtained. These effects were identified in a data-driven manner and represented a distinct dimension of variability in human biology influenced by an external hormone. Thus, the hormonal effects were strong enough to be picked up above all other causes of variability and noise. F14 could therefore be a potentially useful biomarker to assess exposure to EE as well as associated health effects. The association with pulse and the effects on inflammation, adrenal and thyroid function have been individually investigated before^22,23^, but the full depth and width of this signature have not been previously described. EE is a strong synthetic hormone and hence it is not as unexpected that modulation of this pathway could have consequences on the proteome and metabolome as well as body physiology. However, considering that the use of this contraceptive is common, we point out that this will need to be accounted for as a potential bias in all epidemiological and multiomics studies. The EE-inducible signature could otherwise cause strong false interpretations in biomarker studies. For example, serum TFF3 has been reported to be elevated in women with breast cancer^36^ and FETUB is a highly abundant liver-secreted protein sensitive to estrogen and is reported to be increased in type 2 diabetes^37,38^.

Our analysis showed that genetic determinants could explain part of the variability in the multiomics features. This raises the question of whether this type of multi-omics data integration and unsupervised analysis could help to dissect the molecular effects mediating the mechanisms and pathways on how genetics may influence disease onset. Several hints of such observations were seen, such as the associations of IL-6 receptor signalling and abdominal aortic aneurysm.

We acknowledge some limitations of this study. Firstly, the sample size may limit the discovery of weak associations and hence the conclusions should be confirmed in larger cohorts. Nevertheless, we retrieved several known associations, likely resulting from strong underlying effect sizes. For example, we showed that several blood cell measures in our study were under genetic control, in agreement with the high heritability of haematological traits^41^. By extension, it is possible that the associations with cardiometabolic relevance observed in our study also have larger true effect sizes. For example, it was remarkable how key cardiovascular risk genes and drug targets, such as PCSK9 strongly clustered in the network analysis. It has been previously observed that a longitudinal study design substantially increases the power to identify molecular changes in comparison to a groupwise analysis in a population of similar size^4^. Therefore, the presented resource and data analysis could reveal several other relevant interrelationships to be explored further and validated in external data sets. Our work was not a formal interventional study but included longitudinal observations of people undertaking a tailored health intervention. Individual trajectories and multiomics profiles highlighted specific connections when specific cases or subgroups were considered. In addition, in a longitudinal observational study, changes over time may also be caused by other unknown parameters, such as seasonal variability known to impact on immune systems gene expression and cellular composition^42,43^.

Our study illustrated a person-centric data-driven health monitoring that could become part of the future health care practice. This involved integration of clinical, genetic, multi-omics and digital health data as well as unbiased monitoring of longitudinal trends. This study demonstrates the power of combining multi-omics and lifestyle data as well as new opportunities for predicting health trajectories in a holistic, unbiased data-driven manned. This could lead to a re-definition and refinement of the traditional disease-based taxonomy as well as definition of specific transitions between health and disease. This could contribute to preventive approaches via lifestyle and behavioural changes or pharmaceutical interventions.

## Acknowledgements

We thank all the study subjects for their generous participation and commitment. We express our sincerest thanks to Riina Aaltonen, Annette Evokari, Iiro Hietamäki, Anna Seppänen and Paula Vartiainen for their expert assistance. The DHR program was coordinated and managed by the Center for Health and Technology, University of Oulu, Finland. Genotyping was performed by the Institute for Molecular Medicine Finland (FIMM) Technology Centre, HiLIFE, University of Helsinki, Finland.

## Author contributions

F.M, T.J., R.S. and O.K. wrote the paper with contributions from all the authors. F.M. designed and performed the bioinformatics analyses, interpreted the results, prepared the figures and wrote the primary version of the paper. T.J. performed bioinformatic data analysis and interpreted the results. R.S. coordinated the data collection, supervised the experiments, and was responsible for the project management. R.S and O.K. designed the project, with contributions from H.V., H.H. and M.P.H., and provided conceptual advice and support. M.P.H. acted as a coordinator of the DHR consortium. H.S. and Thonas Moritz performed the metabolomics experiments, analysed the data and contributed to the interpretation of the results. F.B. and L.E. performed the microbiome experiments, analysed the data and contributed to the interpretation of the results. C.H., M.N., and P.N performed the autoantibody experiments, analysed the data and contributed to the interpretation of the results. E.W. and S.R. provided the KardioKompassi algorithm. R.M., T.P., M.B. and Timo Miettinen were responsible for data management, database administration, health dashboard and software development. H.L. was the study physician. P.H. performed genotype imputation. Anu Karhu, I.J., K.K., H.V., H.H., Antti Kallonen, M.E., H.S. and M.L. conducted research.

## Methods

### Ethical permit and consent

The study was conducted in line with the Declaration of Helsinki and approved by the Coordinating Ethics Committee of the Helsinki University Hospital, Finland. Each participant provided informed written consent for the study, biobanking of samples, publication and data sharing. Participants were free to drop out any time, but their samples and data were still available for analyses. If a participant withdrew the consent, samples and data were discarded.

### Overview

The observational cohort study was conducted at the Institute for Molecular Medicine Finland (FIMM), HiLIFE, University of Helsinki, Finland, between September 2015 and January 2017. Recruitment was performed in September 2015, and five study visits followed, approximately every four months, (Supplementary Figure 1). Every visit included a health check-up and blood and urine sampling. Before or after every visit, participants collected fecal and saliva samples, filled out a questionnaire, and performed fitness tests. Data on physical activity and sleep was collected from the second visit onward with an activity watch. Key actionable health data were returned to the participants via a web dashboard starting at the second visit. Tailored health and wellness advice and coaching was provided by two personal trainers from the third visit onward. Between visits 4 and 5, participants could compare their data to summary measures calculated with the other participants’ data.

### Eligibility criteria and recruitment

The requirements included: age of 25-64 years at the study start, sufficient computer skills and Internet access via a smartphone compatible with the provided smartwatch; sufficient English language knowledge to be able to understand simple messages and to use health and wellness applications. We excluded individuals with severe diseases (cancer, cardiovascular, debilitating neurological, psychiatric or orthopaedic diseases). However, individuals with risk factors for chronic diseases (e.g., obesity, elevated blood pressure, dyslipidaemias, disturbances in sugar metabolism, mild forms of metabolic syndrome, smoking, moderate drinking or sleep problems) were allowed. Individuals diagnosed with or suspected of having rare monogenic diseases were excluded. In addition, individuals under custody, with special needs or with limited decisional capacity were excluded, as well as people suffering from severe forms of alcoholism and depression. Pregnant women were excluded and women becoming pregnant during the study were no longer asked to provide samples or contribute to the study. Altogether, 645 clients of a private occupational health service (Mehiläinen Töölö, Helsinki, Finland) were invited to participate (Supplementary Figure 1). People interested in participating (n=125) filled out a questionnaire to assess eligibility. A total of 107 volunteers of European descent from the Helsinki metropolitan area were selected to participate and 96 completed the study (Supplementary Table 1).

### Return of personal health data

The ensemble of anthropometrics, clinical laboratory tests, and physiological measurements is called clinical variables in this paper. We returned actionable health data via the Health Dashboard web application, which allowed participants to: explore and compare their data against reference values and the mean values for the other participants, fill out questionnaires and read informational material. The clinical variables and their reference values were returned starting at visit 2. A physician interpreted the clinical data and was available to discuss the results with the participants. The occupational health service communicated any medically relevant finding requiring immediate actions, while the study group communicated non-acute findings. Clinical decisions were made according to the national Current Care Guidelines (https://www.kaypahoito.fi/). Genetic data were returned to participants under the guidance of a clinical geneticist after visit 2. A personal 10-year risk for coronary heart disease (CHD) was communicated using KardioKompassi^44^. The risk model is based on both traditional (sex, age, family history, smoking, systolic blood pressure, total and HDL cholesterol) and hereditary risk factors (approximately 49,000 SNPs). If the risk was >10% the participant was referred to a doctor. We also communicated the risk for low serum 25(OH)D concentration after visit 4 (Sallinen et al., The Journal of Nutrition, in press). We returned visualizations of questionnaire data on diet, physical activity, sitting, sleeping, subjective health status, and mental wellbeing after visit 4. We also determined a self-reported obstructive sleep apnea risk score^45^. Participants were able to monitor their physical activity and sleep data using the Withings Health Mate app starting at visit 2.

### Coaching and group meetings

Two personal trainers (PTs) coached participants from visit 3 onward. Coaching included nine 30-minute private meetings (three face-to-face and six remote), email/phone support, as well as group meetings. The PTs were accredited by the European Health and Fitness Association. No structured, evidence-based coaching protocol was used, but personal actionable possibilities were defined to help participants change their behaviour and improve their health. With the help of a physician and a nutritionist, the PTs translated and customized actionable possibilities to specific recommendations. The PTs had access to participants’ age, clinical variables, fitness test data, and information about their occupation, exercise, and diet. Tailored health advice and coaching were based on each participant’s behaviour, health risks, lifestyle, and goals. One to three relevant, personalized and actionable opportunities were offered to guide and motivate participants to make lifestyle changes to optimize wellness and health and delay predicted pathologies. Personalized advice usually focused on diet, exercise, mental wellbeing, or stress and time management. If necessary, a physician limited exercise. Twenty-one informational group meetings were organized during the study, where participants could interact with a physician, a nutritionist, a nurse, PTs, and other experts. Participation in these meetings was voluntary except for the first, during which a physician and a geneticist interpreted the clinical variables data as well as the CHD risk prediction.

### Experimental procedures

#### Health check-up and clinical variables

Health check-ups occurred on weekdays between 7:30 and noon. Body weight, height, waist and hip circumferences, blood pressure, and pulse were measured using standard procedures. Participants were asked about medications, dietary supplements, and the use of health care services. Blood and urine samples were collected at every visit. Fasting (≥8 h) blood samples were collected, and clinical chemistry assays were performed on blood, plasma or serum, at the diagnostic laboratory of Mehiläinen Töölö (Helsinki, Finland), United Medix Laboratories Ltd (Helsinki, Finland) and HUS Diagnostic Center (Helsinki, Finland). Urine samples (≥2 h without urinating) were collected using standard procedures. A dipstick test was performed on the urine samples at the diagnostic laboratory of Mehiläinen Töölö (Helsinki, Finland). Aliquots of blood, plasma, serum and urine were frozen at −20°C and stored in liquid nitrogen for biobanking.

### Questionnaire

Before every visit, participants filled out an online questionnaire, including approximately 150-190 questions, depending on the visit, and covering the following: personal information; sociodemographic and socioeconomic characteristics; familial and individual disease history; functional capacity and health; mental wellbeing; physical activity and exercise habits; diet (including food frequency questionnaire, FFQ); smoking; alcohol consumption; sleep; occupational health; personality traits; attitudes and expectations towards lifestyle changes; monitoring own health and wellbeing; expectations towards health technology. Healthy Food Intake Index is a food-based diet quality index, calculated from the FFQ data and adapting an available method^46^.

### Fitness tests

After every visit, participants were instructed to perform lower-body (squats, repetitions/30s), abdominal (sit-ups, repetitions/30s) and upper-body (push-ups, max number of repetitions) muscular fitness, balance and mobility tests. Participants executed the tests and uploaded the results to the Health Dashboard.

### Activity and sleep monitoring

After the second visit, participants were equipped with the Withings Activité Pop smartwatch, connected to the Withings Health Mate app, to measure physical activity, energy expenditure and sleep. Participants were requested to wear the watch until the end of the study. Wellness Warehouse Engine^47^ embedded in the Health Dashboard was used to authorize and provide the research group access to participants’ data. Only aggregated daily activity and sleep data were exported and included in this study.

### Salivary cortisol

Participants collected saliva samples within two weeks of every visit, using Cortisol-Salivette® with a synthetic swab (Sarstedt). Four samples (awakening, 15 min and 30 min after wake up, bedtime) were collected on a typical weekday. Time and mood at sampling were recorded. Samples were stored at +4°C until aliquoting and storage at −20°C. Salivary cortisol levels were determined at the University of Trier (Trier, Germany) using a DELFIA immunoassay^48^. The stress scores derived from the measurements included: AUCi, AUCg (according to^49^), awakening, evening, peak and Delta (peak - ground) cortisol levels.

### DNA extraction and genotyping

DNA was extracted from whole blood with Chemagic MSM1 (PerkinElmer). Genotyping was performed at the FIMM Technology Centre (HiLIFE, University of Helsinki, Finland) using InfiniumCoreExome-24 v1.0 DNA Analysis Kit, iScan system and standard reagent and protocols (Illumina). Genotypes were pre-phased with ShapeIT2 and imputed with IMPUTE2. Two pre-phased reference panels were used (--merge_ref_panels): 1000G Phase 1 and a Finnish low-coverage WGS imputation reference panel, composed of 1,941 whole-genome sequences (SISu project). Imputation resulted in 30.3 million quality-filtered variants (R^2^>0.3 and missingness<0.05).

### GCMS and LCMS

GCMS and LCMS experiments were performed at the Swedish Metabolomics Center (Umeå). 100 μl of plasma were processed as described^50^, with the following details: extraction with 900 μl of 90% v/v methanol, containing internal standards for both GCMS and LCMS, at 30 Hz for 2 minutes; protein precipitation at +4 °C; centrifugation at +4 °C, 14 000 rpm, 10 minutes. 50 or 200 μl of supernatant were evaporated to dryness in a speed-vac concentrator, for GCMS or LCMS analysis respectively, and stored at −80 °C. Quality control (QC) samples were created by pooling supernatants. MSMS analysis (LCMS) was run on the QC samples for identification purposes. Sample batches were created according to a randomized run order. For GCMS, derivatization and analysis were performed as described previously^50^, with the following modifications: 0.5 μL of derivatized sample; splitless injection with a L-PAL3 autosampler (CTC Analytics AG); 7890B gas chromatograph (Agilent Technologies); chemically bonded 0.18 μm Rxi-5 Sil MS stationary phase (Restek Corporation); column temperature increased from 70 to to 320 °C at 40 °C/min; Pegasus BT TOFMS (Leco Corporation); solvent delay of 150 s; detector voltage 1800-2300 V. Unprocessed MS-files were exported from the ChromaTOF software in NetCDF format to MATLAB R2016a (Mathworks), where processing occurred, including baseline correction, chromatogram alignment, data compression and Multivariate Curve Resolution. The extracted mass spectra were identified by comparisons of their retention index and mass spectra with known libraries using the NIST MS 2.0 software^51^. Annotation was based on reverse and forward searches in the library. LCMS experiments were performed as described^52^ and all data processing was performed using the Masshunter Profinder version B.08.00 (Agilent Technologies). The processing was performed both in a target and an untargeted fashion. For target processing, a predefined list of metabolites commonly found in plasma and serum were searched for using the Batch Targeted feature extraction in Masshunter Profinder. An-in-house LCMS library, built up by authentic standards run on the same system with the same chromatographic and MS settings, was used for the targeted processing. The identification of the metabolites was based on MS, MSMS and retention time information.

### Autoantibody bead arrays

Autoantibody profiling on antigen bead arrays was performed as previously described^27^, at the Autoimmunity and Serology Profiling facility at SciLifeLab (Stockholm).

### PEA

Plasma proteins were quantified using Proximity Extension Assay (PEA) at Olink Bioscience (Uppsala), using 6 panels: Cardiometabolic, CVD II, CVD III, Inflammation I, Metabolism and Oncology II. Each panel assayed 92 proteins using a matched pair of antibodies coupled to oligonucleotides, which form an amplicon by proximity extension and can be quantified by real-time PCR. The data were normalized against an extension control and an interplate control and expressed as Normalized Protein eXpression (NPX) values, which represent an arbitrary relative quantification unit on log2 scale. The NPX values below the limit of detection (LOD) were considered missing and their value was replaced with the LOD value for each assay.

### Microbiome analysis

Participants collected faecal samples within two weeks of every visit. Samples were collected in precooled vials placed in styrofoam boxes with ice gel packs and then stored at +4°C for a maximum of one day until aliquoting, freezing at −20°C, and storage in liquid nitrogen without additives. Thawed faecal samples were spun-down and DNA was extracted, with 50 ng of DNA submitted to PCR amplification as described^53^, using 341f and 805r primers (CCTACGGGNGGCWGCAG and GTGBCAGCMGCCGCGGTAA) for the V3−V4 regions of 16S rRNA^54^. Sequencing was done on an Illumina MiSeq with 2×250 bp reads. After quality trimming with Cutadapt, an ASV (amplified sequence variants) table was generated using the DADA2 pipeline^55^, including the following steps: filtering and trimming, learning of error rates, dereplication, sample inference, read pairs merging, removal of chimeras and taxonomy assignment using SILVA v128 database.

### Bioinformatic analyses

#### Longitudinal analysis

We applied Generalized Estimating Equations (GEE) to the quantitative clinical variables to investigate the longitudinal changes while controlling for sex and age. As data feedback and coaching started respectively from visit two and three, we excluded the first visit and considered the second visit as baseline. Four time points (0-4) were then considered and we aimed at extracting the average change per visit, assuming the same change in the values between any two successive visits. Furthermore, for each variable, individuals were classified as Out-Of-Range at Baseline (OORB) if the values were outside the reference range (Supplementary Table 2) at baseline. We assumed that the longitudinal changes could be distinct for individuals classified as ORRB. In other words, we allowed the effect of coaching and data feedback to be different in the two groups by introducing an interaction term. If the OORB group included at least five individuals, then the variable was modelled as:

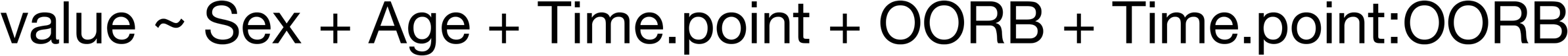

Otherwise, a model with no groups of individuals was considered:

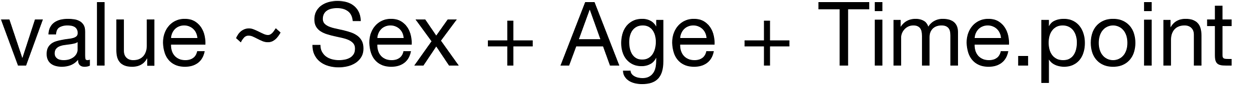

GEE models were fitted using the geepack library and exchangeable correlation structure. We extracted the effects, their standard errors and p-values using the esticon function in the doBy library. If no interaction was specified, the coefficient for Time.point represented the estimated change for all the individuals, expressed as the change occurring between two successive visits. If the interaction was included, we considered the change occurring in the ORRB group between two successive visits, and we extracted the effect as the linear combination of the coefficients of Time.point and OORB. P-values were adjusted with the Benjamini-Hochberg method. Model prediction of the variable values was visualized by stratifying for sex and OORB group. The 95% confidence levels were estimated by bootstrapping with 1000 replications.

### Dimensionality reduction

We selected the quantitative clinical variables and imputed missing values using the `imputePCA` function in the missMDA package. Principal Component Analysis (PCA) was performed with the `PCA` function in the FactoMineR package on scaled data. Variable contribution to a given PC was defined as the ratio between the squared cosine of a variable and the sum of the squared cosines for all the variables for that component.

### Data preprocessing

#### Activity and sleep

We obtained summary measures for the activity and sleep data by calculating average values for the different variables in the previous 30 days before each visit. We considered only samples for which at least 10 days were included in the calculation.

#### GCMS

The relative quantities (RQs) were log2-transformed and missing values were imputed with the KNN algorithm. Data were normalized using the RUV4 method in the ruv R library^56^. `RUV4` was applied to remove sources of technical variability, including batch effect and the signal drift over time. Briefly, we used the internal standards (IS) as negative controls, k=4 and a design matrix with individuals and visits as covariates, in order to estimate the matrix W of unwanted factors and their respective α coefficients. The number of factors to remove (k) was chosen with the `getK` routine, but was ensured not to exceed 1/3 of the number of the IS. The normalized relative concentrations were obtained by subtracting the effect of the W components from the RQs. The IS values were inspected before and after normalization to ensure a removal of the signal drift over time and a reduction of the Coefficient of Variation (CV). Estimation of the Intraclass Correlation Coefficient (ICC) and PCA plots were also evaluated to inspect the performance of the normalization method.

#### LCMS

The negative and positive modes from LCMS experiments were processed separately. Unidentified metabolite peaks and metabolites with more than 25% of missing data were removed. RQs were log2-transformed and missing values were imputed with KNN. `RUV4` adjustment and inspection of the results were performed as described above with k=2 or k=1 for LCMS-neg and LCMS-pos data, respectively.

#### Autoantibodies

We considered: continuous Median Fluorescence Intensity (MFI), discrete binned and binary (0=undetected, 1=detected) data (see ^27^). MFI values were normalized with the Probabilistic Quotient Normalization^57^, with a median reference profile, and then log2-transformed.

#### 16S rRNAseq

The ASV count table represents an analogue of the traditional Operational Taxonomic Unit table. We inspected the taxa prevalence (the number of samples in which a taxon has nonzero counts) and the total abundance (sum of the counts in all samples) and the abundance distribution. Using phyloseq R library, we agglomerated the counts to the genus level, to reduce functional redundancy. The genus-level data were filtered to remove taxa with missing phylum annotation, prevalence ≤2 and total abundance ≤10. We inspected the ordination of the samples using several distance metrics. We then converted the genus-level count to a DESeq2 dataset and used the Variance Stabilizing Transformation (VST), which normalizes with respect to the library size and gives a matrix of approximately homoscedastic values that can be used for downstream analyses. As a major source of batch effect was represented by the experimental plate, we verified that the VST conversion also reduced the library size effect by visualizing the sample ordination, using MDS with Bray-Curtis distance for the genus-level counts and Euclidean distance for the VST values. We also formally tested the reduction of the batch effect, by looking at the association between the first two axes of the sample ordination plots with the plate ID. The estimation of the α-diversity for each sample was done using the original unfiltered counts.

### Integration of multiomics experiments

A MultiAssayExperiment object was assembled with all the continuous clinical and omics data, namely RUV4-nomalized data for GCMS, LCMS-neg and LCMS-pos, log2MFI for autoantibodies, NPX values for PEA assays and VST-transformed values for 16S counts. Data filtering to remove unwanted features included: removal of internal standards from GCMS and LCMS; removal of proteins NPX values < LOD (i.e. missing data) in more than 25% of the samples; removal of FS, CCL22 and BDNF (technical issues, Olink communication); removal of autoantibodies with background fluorescence in all samples (scored reactivity values ≤0.5); removal of bacterial taxa with a prevalence ≤30%. We used the filtered object for all the downstream analyses. The resulting dataset included 136 GCMS metabolites, 104 LCMS-neg metabolites, 163 LCMS-pos metabolites, 174 autoantibodies, 501 proteins and 85 bacterial taxa (Supplementary table 2).

### Variance partition analysis

For each omics type, between- and within-individual variation was inspected with a distance-based method and with variance partitioning analysis. For the distance method, Euclidean distance between all pairs of samples was calculated. Then, the within-individual distance was calculated as the median value of the distance values of the sample pairs from the same individual. This value was plotted together with the distribution of the remaining distance values, i.e. the distances between one individual and the rest of the individuals. For the variance partitioning method, a linear mixed model was fitted with lme4 R library, independently to each feature with nonzero variance. We included the individual as a random intercept and age, sex and time point (0-4) as fixed effects. The fixed effect variance, random intercept variance and residual variance components were extracted, and the relative fraction calculated. The random intercept variance represented the between-individual variance, while we interpreted the residual variance as the within-individual variance not accounted by the fixed effects.

### Multi-Omics Factor Analysis

Multi-Omics Factor Analysis v2 (MOFA+) provides a set of factors that capture biological and technical sources of variability^11,12^. It infers the axes of heterogeneity that are shared across multiple modalities and those specific to individual data modalities. We used the “intercept_factor” branch of the code repository (https://github.com/bioFAM/MOFA2/tree/intercept_factor). For training, we selected the samples that had a measurement in all the omics layers (n=359). We considered a Gaussian likelihood for the continuous measurements for all the layers, except for autoantibodies, which were considered in their binary form (0=undetected, 1=detected) with Bernoulli likelihood. Data were not scaled. We trained 10 alternative models in Python with a random seed, starting from 20 factors and the following options: iter=”10000”, convergence_mode=”medium”, dropR2=”0.02”. The model with the best value of the Evidence Lower Bound (ELBO) was selected. We checked the robustness of the learned factors by inspecting the Pearson correlation between the factors obtained by all the runs (Supplementary Figure 6). We then computed the fraction of the total variance explained by each factor and the fraction of variance explained by each factor in each layer. We inspected the factor loadings to understand the contribution of the original features. A feature with a higher absolute loading has a higher weight on the factor, but because the loadings are not directly comparable, their scaled values were used for visualization. The sign of the loading defines a direct (positive) or inverse (negative) proportionality with the corresponding factor, and features with similar loadings contribute similarly to the factor. We inspected the factor values and to visualize and cluster the samples on the reduced space.

### Association between factors and phenotypic variables

In order to test the association between the MOFA+ factors and the sample characteristics, we gathered a collection of phenotypic information including: clinical variables; questionnaire data (family history, exercise and physical activity, functional capacity and health, mental health, diet, smoking, alcohol use and sleep habits); Healthy Food Intake Index; KardioKompassi risk; sleep apnea score; stress scores; fitness tests; sleep and activity summaries from the wearables. For each combination of factor and phenotypic variables, we fitted a linear model with the factor values as outcome and the phenotypic variable as predictor. Reported R^2^ values were derived from these models. For the categorical variables, the predictor was considered an ordered factor if the factor levels corresponded to ordered levels and the variable could be considered as a qualitative or semi-quantitative ordinal variable. A False Discovery Rate (FDR) was calculated using the Benjamini-Hochberg method. The candidate associations for further screening were obtained at FDR<0.05, while the associations reported in Figure 3 satisfy a FDR<0.001 threshold, in order to better control for false positives arising from the multiple hypotheses tested. The selected associations were refined with linear mixed models as implemented in lme4 library. We modelled the LF values with the individual as a random intercept and age and sex as fixed effect. We refer to these models as the covariate-adjusted models. P-values for these models were obtained with the “Type II ANOVA” as implemented in the Anova function in the car library, with a Kenward-Roger F test.

### Population stratification

The DHR and the 1000 Genome phase 3 genotypes were merged. Analyses were done with PLINK v1.9. Only autosomal biallelic SNPs with genotyping rate >95% and MAF>5%. were considered. A set SNPs in approximate Linkage Disequilibrium was obtained with the option --indep-pairwise 50 5 0.2 and PCA was performed. To predict the ethnicity, a Linear Discriminant Analysis model (`lda` in MASS) was trained in R with the PCA scores of the 1000 Genomes only and tested on the DHR samples.

### GWAS summary scores

We considered 146 traits from the GWAS catalog v1.0.1 (https://www.ebi.ac.uk/gwas/) for selected ontology categories: body measurement, cardiovascular disease, cardiovascular measurement, hematological measurement, inflammatory measurement, lipid or lipoprotein measurement, liver enzyme measurement, metabolic disorder and other (fasting blood glucose, vitamin D levels and thyroid stimulating hormone). For each trait, only the studies meeting the following criteria were considered: at least one SNP with p-value<1E-08, sample size >5000, at least 10 SNPs reported. The study with the largest sample size was considered for the traits with multiple associated studies. For each biallelic SNPs, the reported effect size of the risk allele was considered as a weight. Summary scores were computed by multiplying the imputed genotype dosage of each risk allele times its respective weight and summing across all SNPs.

### Between- and within-individual cross correlation network

We considered the continuous measurements for all the omics layers, together with a collection of measurements including: GWAS summary scores; gut microbiome alpha diversity; clinical variables; questionnaire data; Healthy Food Intake Index; KardioKompassi risk; sleep apnea score; stress scores; fitness tests; sleep and activity summaries. The resulting dataset was preprocessed by regressing out the effect of sex and then the effect of age for each variable, only if significantly associated. For the between-individual cross correlation network calculation, the observations were grouped by individual and averaged, obtaining 96 independent observations for each of the 1394 variables. Spearman correlation was calculated for each pair of variables, resulting in 731318 nonmissing estimated ρ coefficients, p-values and Benjamini-Hochberg adjusted p-values (FDR), for the pairs of features of different types (activity and sleep, autoantibody, clinical variables, fitness test, genetic, HFII, KardioKompassi, metabolite, microbial, protein, sleep apnea score, stress score). For the within-individual cross-correlation network calculation, the GWAS scores and the autoantibody data were excluded, resulting in 1058 variables, 96 individuals and 5 time points. We used rmcorr library^58^ to estimate the common within-individual association for grouped values measured at the five visits. `rmcorr` estimates a common regression slope representing the association shared among individuals and provides the best linear fit for each individual using parallel regression lines with different intercepts. The repeated-measures correlation coefficient (r_rm_) is similar to the Pearson correlation coefficient and measures the strength of a linear association but, unlike Pearson correlation, it takes into account non-independence between the measures. Hence, we estimated 347640 nonmissing r_rm_, p-values and Benjamini-Hochberg adjusted p-values (FDR), for the pairs of features of different types and with at least 100 nonmissing pairs of values. For downstream analysis, we selected the associations satisfying the condition FDR<0.05 and coefficient>0.3 (ρ or r_rm_) and generated two annotated correlation networks. Communities were extracted with the Louvain method, resulting in modularity values of 0.70 and 0.69 for the between- and within-individuals correlation network respectively.

---

**Supplementary Table 1.**
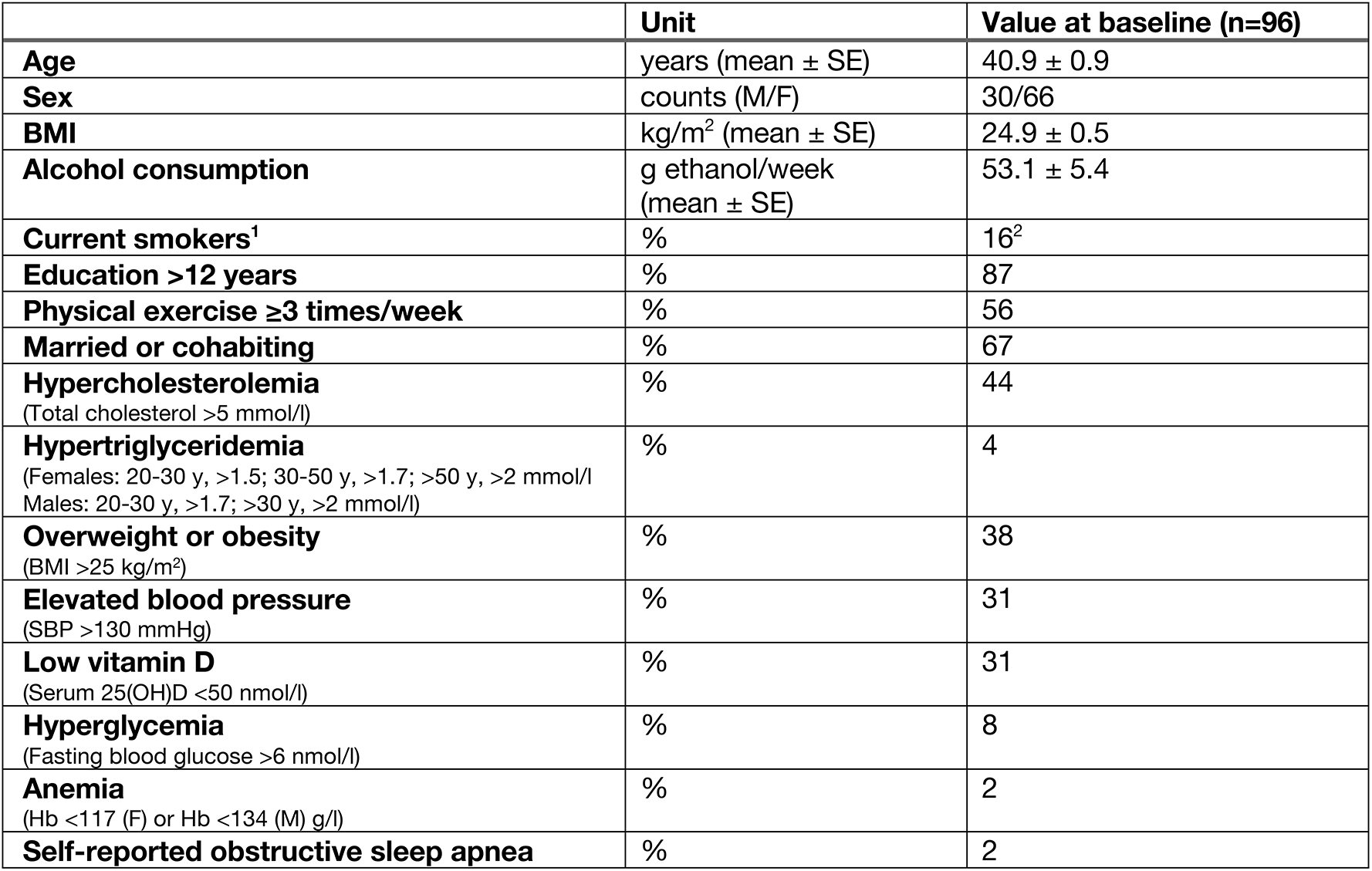
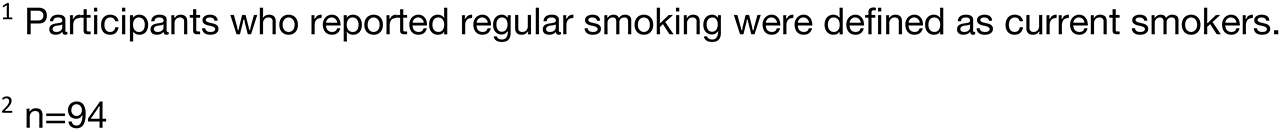
Characteristics of the study cohort.

|  | **Unit** | **Value at baseline (n=96)** |
| --- | --- | --- |
| **Age** | years (mean ± SE) | 40.9 ± 0.9 |
| **Sex** | counts (M/F) | 30/66 |
| **BMI** | kg/m<sup>2</sup> (mean ± SE) | 24.9 ± 0.5 |
| **Alcohol consumption** | g ethanol/week (mean ± SE) | 53.1 ± 5.4 |
| **Current smokers<sup>1</sup>** | % | 16<sup>2</sup> |
| **Education >12 years** | % | 87 |
| **Physical exercise ≥3 times/week** | % | 56 |
| **Married or cohabiting** | % | 67 |
| **Hypercholesterolemia**<br>(Total cholesterol >5 mmol/l) | % | 44 |
| **Hypertriglyceridemia**<br>(Females: 20-30 y, >1.5; 30-50 y, >1.7; >50 y, >2 mmol/l<br>Males: 20-30 y, >1.7; >30 y, >2 mmol/l) | % | 4 |
| **Overweight or obesity**<br>(BMI >25 kg/m<sup>2</sup>) | % | 38 |
| **Elevated blood pressure**<br>(SBP >130 mmHg) | % | 31 |
| **Low vitamin D**<br>(Serum 25(OH)D <50 nmol/l) | % | 31 |
| **Hyperglycemia**<br>(Fasting blood glucose >6 nmol/l) | % | 8 |
| **Anemia**<br>(Hb <117 (F) or Hb <134 (M) g/l) | % | 2 |
| **Self-reported obstructive sleep apnea** | % | 2 |

<sup>1</sup> Participants who reported regular smoking were defined as current smokers.
<sup>2</sup> n=94

**Supplementary Table 2. List of all the included features and their annotation.**

(see separate Excel file)

**Supplementary Table 3.** Results from the GEE model.

|  | **Estimate** | **Units** | **CI** | **P-value** | **FDR** | **Type** | **# of OORB** |
| --- | --- | --- | --- | --- | --- | --- | --- |
| **Hip circumference** | -0.667 | cm | -0.957 – -0.377 | 6.34E-06 | 8.87E-05 | All | 1 |
| **Serum 25(OH)D** | 4.072 | nmol/l | 2.336 – 5.808 | 4.29E-06 | 8.87E-05 | OORB | 24 |
| **Systolic blood pressure** | -3.078 | mmHg | -4.469 – -1.688 | 1.43E-05 | 1.33E-04 | OORB | 27 |
| **Diastolic blood pressure** | -0.947 | mmHg | -1.443 – -0.451 | 1.81E-04 | 1.01E-03 | OORB | 22 |
| **Pulse** | -3.214 | bpm | -4.876 – -1.551 | 1.52E-04 | 1.01E-03 | OORB | 9 |
| **Total/HDL cholesterol ratio** | -0.405 |  | -0.66 – -0.150 | 1.88E-03 | 8.77E-03 | OORB | 7 |
| **LDL cholesterol** | -0.085 | mmol/l | -0.139 – -0.030 | 2.21E-03 | 8.83E-03 | OORB | 29 |
| **ApoB/ApoA1 ratio** | -0.014 |  | -0.024 – -0.005 | 3.89E-03 | 1.36E-02 | OORB | 35 |
| **Cholesterol** | -0.084 | mmol/l | -0.145 – -0.024 | 5.83E-03 | 1.81E-02 | OORB | 36 |

**Supplementary Table 4. Associations between MOFA factor values and phenotypic variables.**

(see separate Excel file)

**Supplementary Table 5. Node and Edge tables for the Between- and Within-Individual Networks.**

(see separate Excel file)

