## Supplementary figures for "Multiomics and digital monitoring during lifestyle changes reveal independent dimensions of human biology and health"

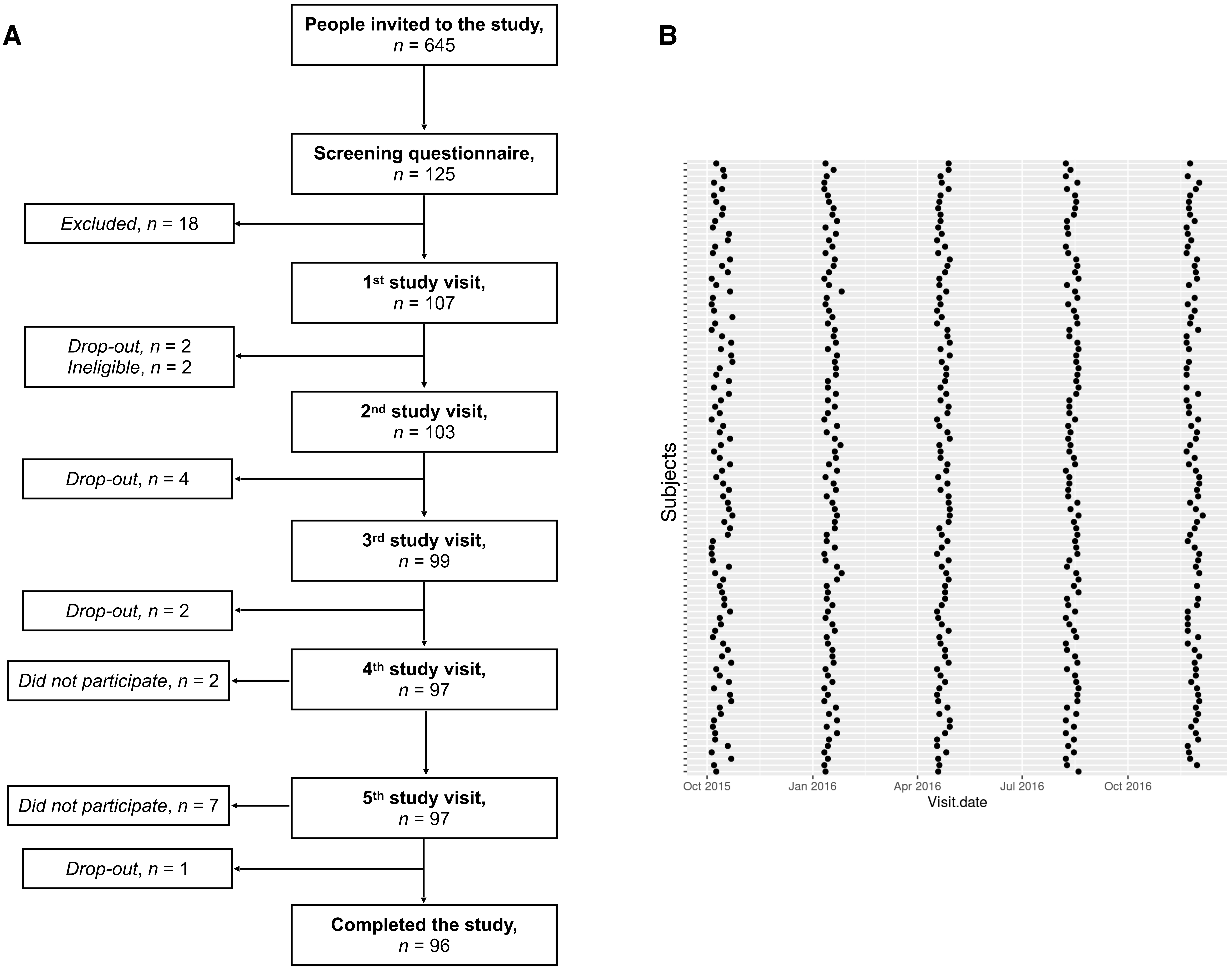


**Supplementary figure 1: A**. Study flowchart showing the number of individuals from the recruitment to the en of the study. **B**. The visit date for each individual is shown. Every row corresponds to an individual and the dots to the corresponding five study visits.

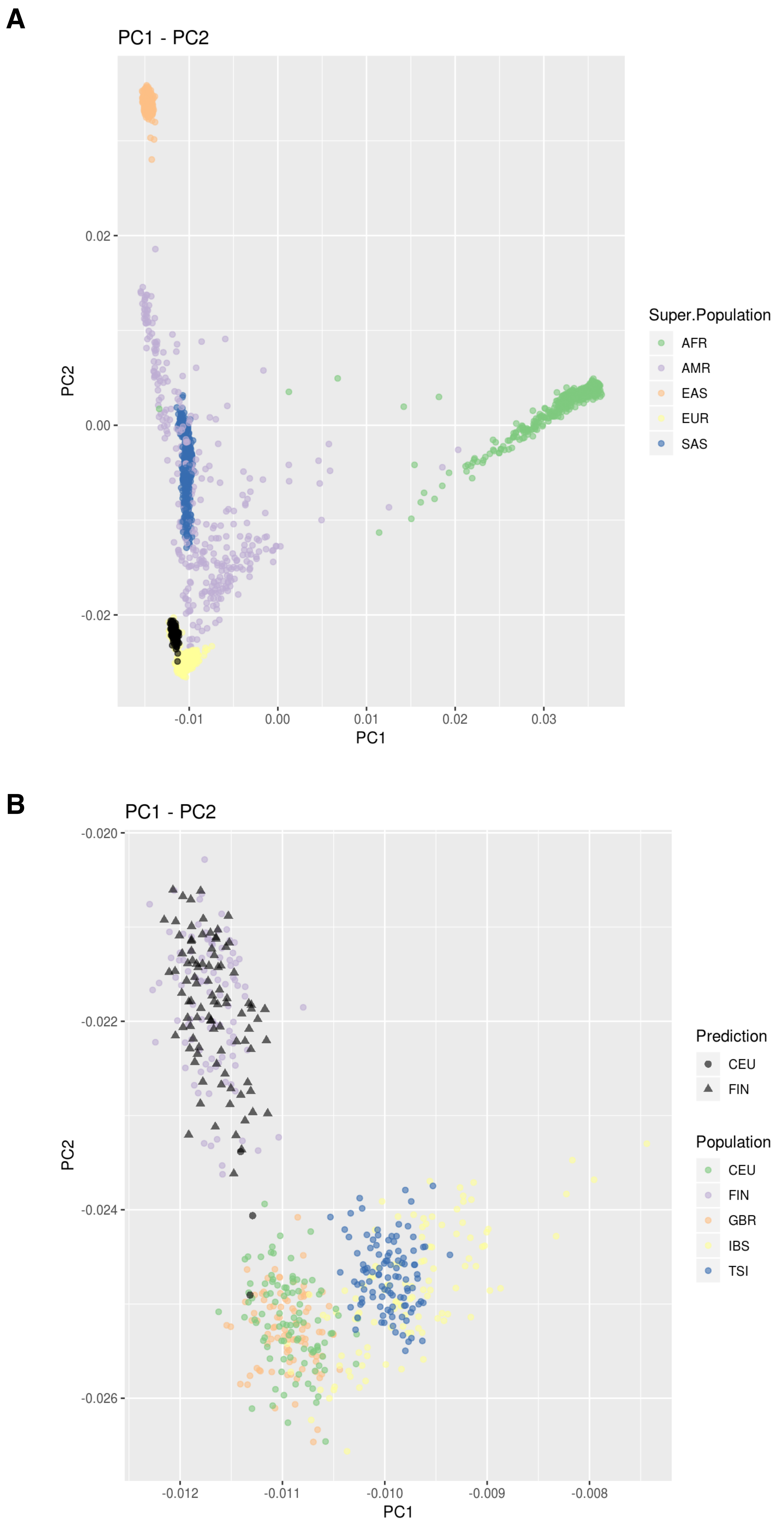


**Supplementary figure 2:** Population stratification plots obtained with PCA analysis using a LD-pruned biallelic set of SNPs. The DHR individuals are shown in black while the other individual from the 1000 Genome phase 3 project are shown in different colors, according to their respective population. **A.** All 1000G samples together with the DHR individuals. **B**. Only the individuals from the EUR super-population are shown with colored points. The DHR individuals are shown in black, while the symbol denotes the predicted population according to a Linear Discriminant Analysis model

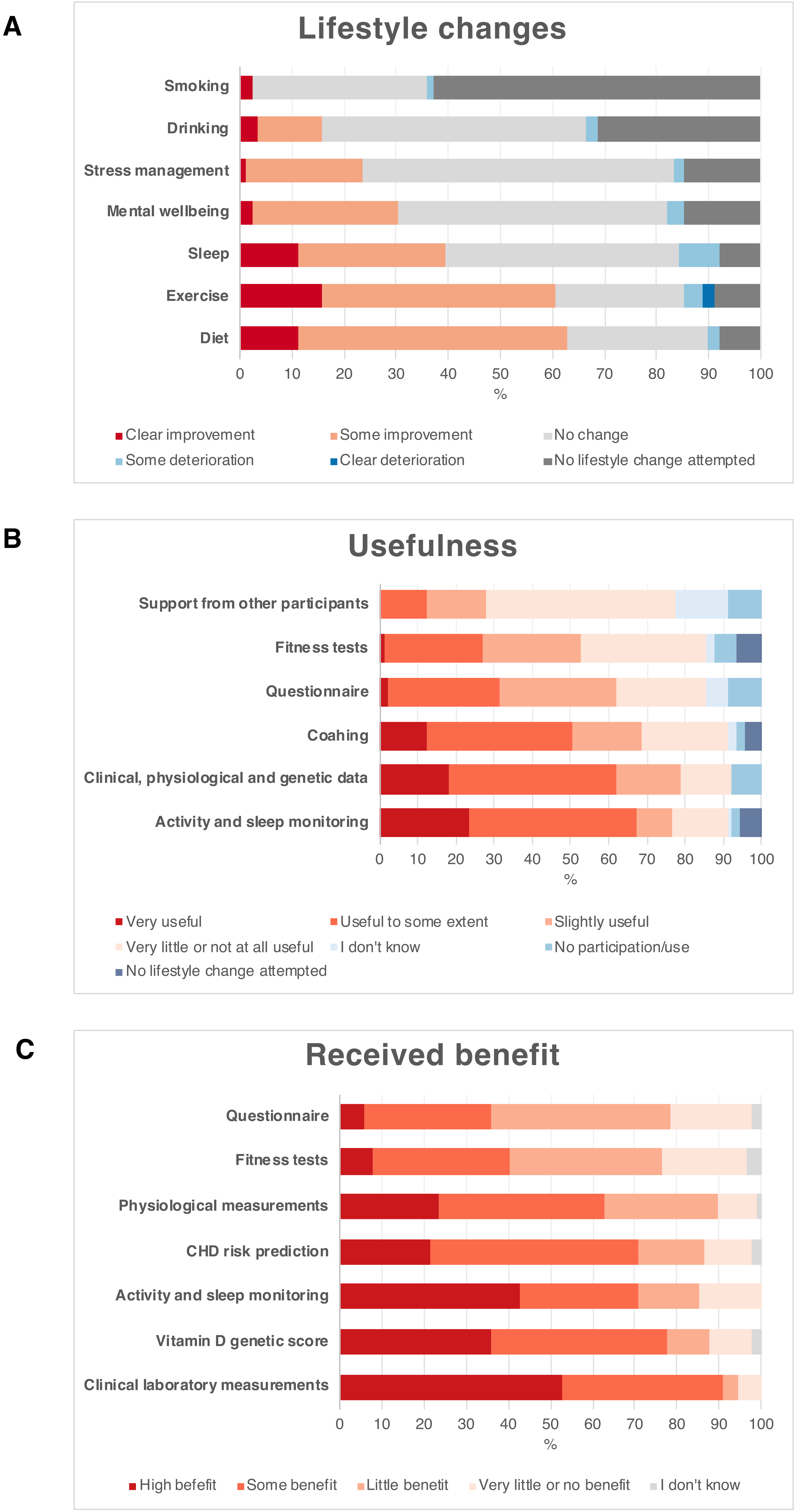


**Supplementary figure 3. A.** Perceived lifestyle changes. Question: "Do you feel that a change has occurred in the following factors during the study?". **B.** Usefulness of the data to make lifestyle changes. Question: "Were the following useful to make lifestyle changes?" **C.** Received benefit. Question: "Did you benefit from the following results/information that you received during the study?"

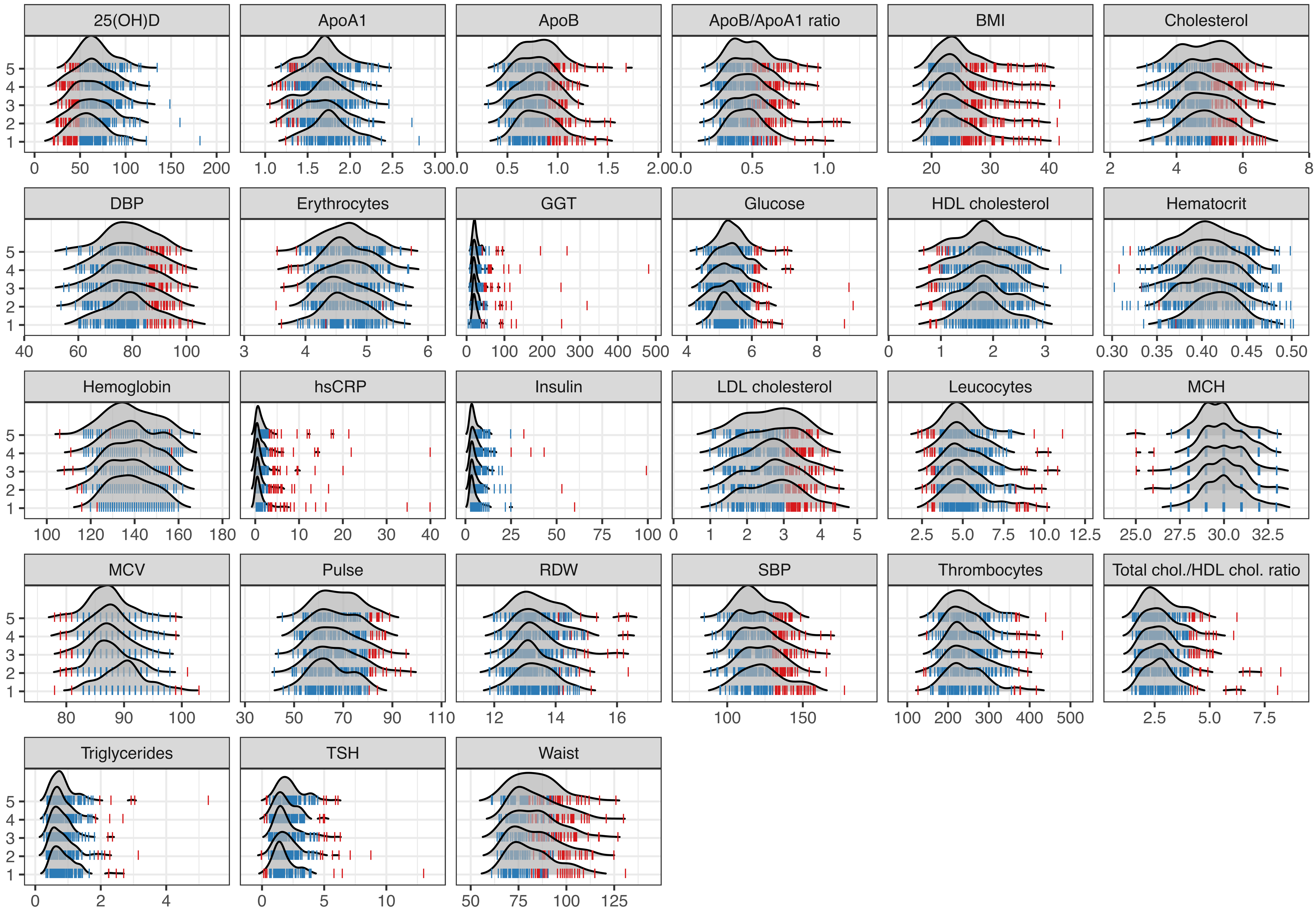


**Supplementary figure 4:** Distribution of the values of the clinical variables. Density distributions were obtained for each visit and variable and shown separately. Vertical lines indicate individual values and are coloured according to their Out-Of-Range status (blue: within range, Red: out of range).

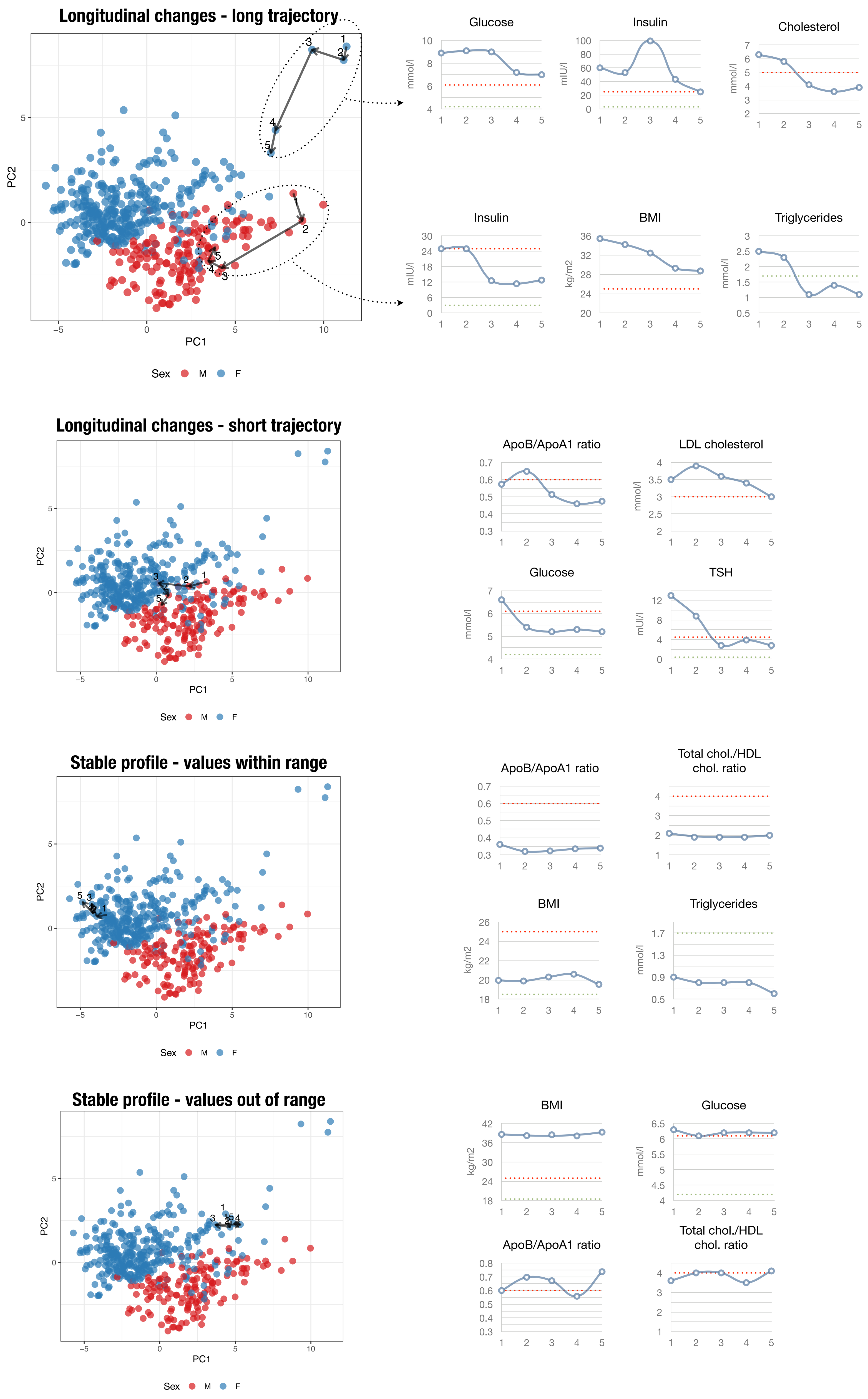

Longitudinal changes - long trajectory
PC1
PC2
Sex M F
Glucose mmol/l
Insulin mIU/l
Cholesterol mmol/l
Insulin mIU/l
BMI kg/m2
Triglycerides mmol/l
Longitudinal changes - short trajectory
PC1
PC2
Sex M F
ApoB/ApoA1 ratio
LDL cholesterol mmol/l
Glucose mmol/l
TSH mIU/l
Stable profile - values within range
PC1
PC2
Sex M F
ApoB/ApoA1 ratio
Total chol./HDL chol. ratio
BMI kg/m2
Triglycerides mmol/l
Stable profile - values out of range
PC1
PC2
Sex M F
BMI kg/m2
Glucose mmol/l
ApoB/ApoA1 ratio
Total chol./HDL chol. ratio

**Supplementary figure 5 (previous page):** Extended main figure 2. **A.** Principal components plot of the individuals obtained using all the numerical clinical variables. The trajectories for five individuals across successive study visits are shown with arrows, along with the corresponding values for selected clinical variables.

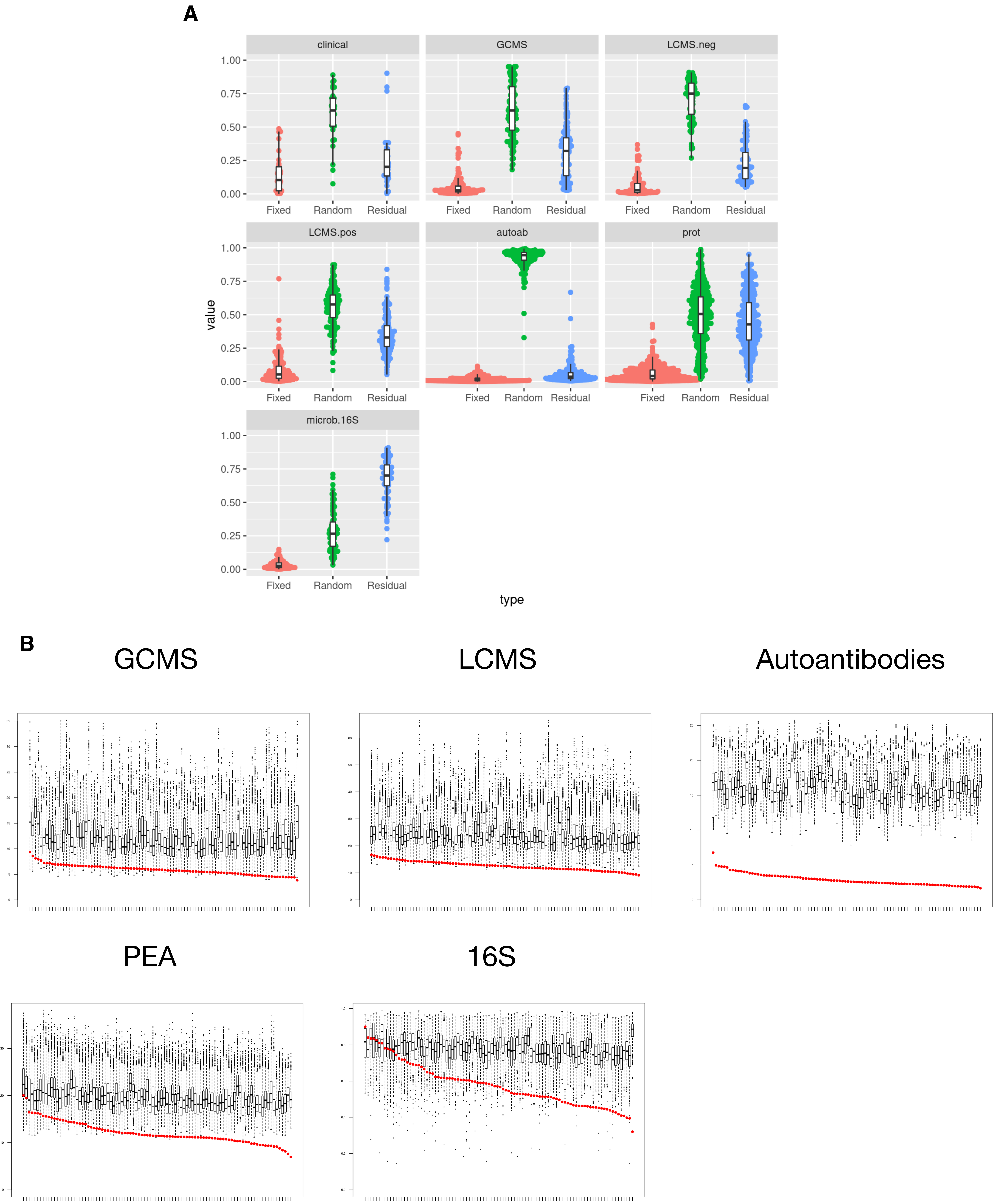


**Supplementary figure 6:** Variance partitioning analysis. **A.** A linear mixed model was fitted independently to each feature. We included the individual as a random intercept and age, sex, time point and phenotypic subgroup as fixed effects. The fixed effect variance, random intercept variance and residual variance components were extracted and the relative fraction plotted. **B.** Each column represents an individual and show the distribution of the Euclidean distances between each individual and the rest of the individuals. A red dot indicates the median value of the distances obtained with the pairs of samples from the same individual (whitin-individual distance).

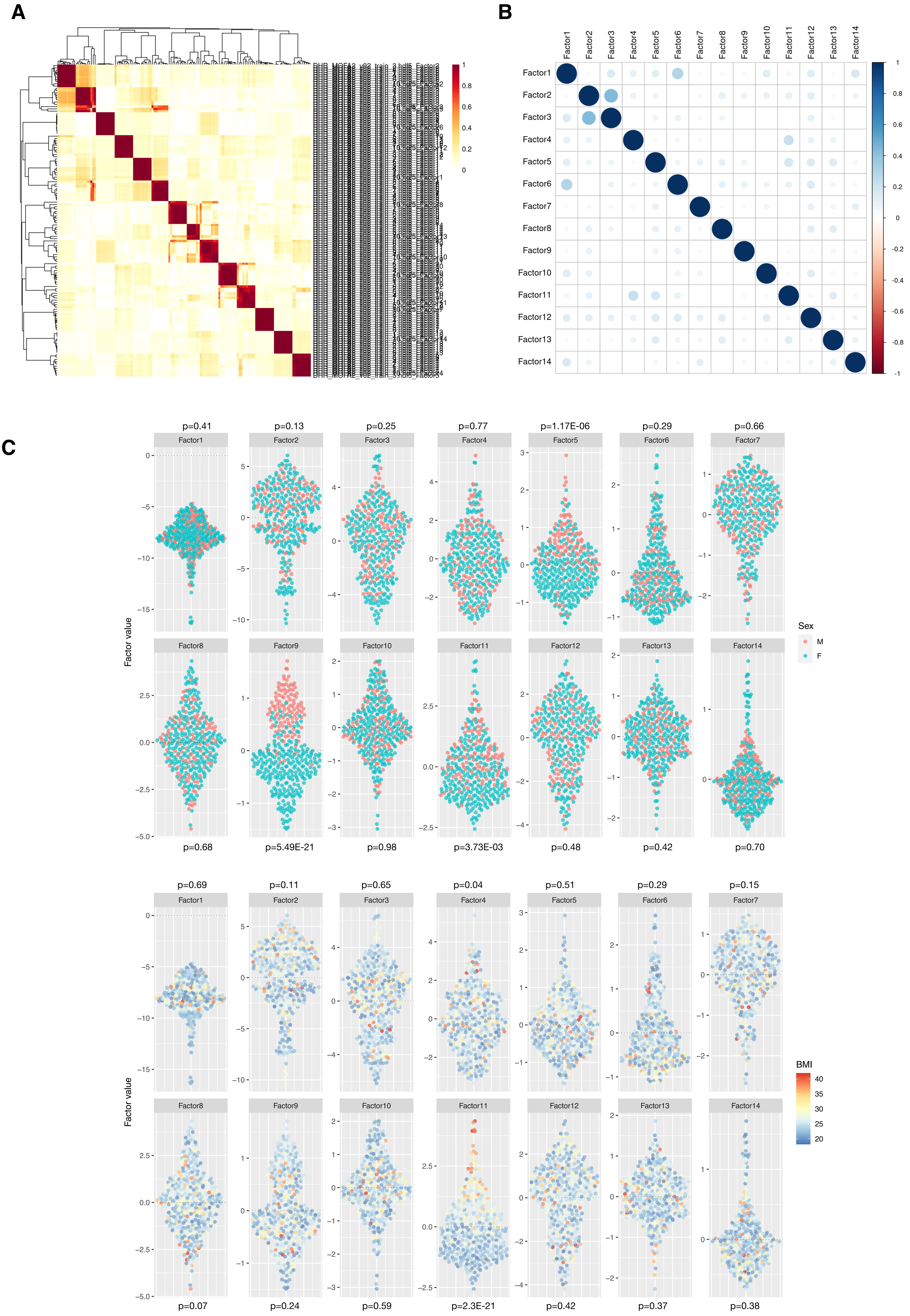

A
B
C
Factor1
Factor2
Factor3
Factor4
Factor5
Factor6
Factor7
Factor8
Factor9
Factor10
Factor11
Factor12
Factor13
Factor14
p=0.41
p=0.13
p=0.25
p=0.77
p=1.17E-06
p=0.29
p=0.66
p=0.68
p=5.49E-21
p=0.98
p=3.73E-03
p=0.48
p=0.42
p=0.70
Sex
M
F
Factor value
p=0.69
p=0.11
p=0.65
p=0.04
p=0.51
p=0.29
p=0.15
p=0.07
p=0.24
p=0.59
p=2.3E-21
p=0.42
p=0.37
p=0.38
BMI
40
35
30
25
20

**Supplementary figure 7 (previous page)**: **A:** Ten different models were trained with random initialization and then the consistency of the learned factors across multiple runs was assessed with the absolute Pearson correlation coefficients between pairs of factors. **B:** The model with the highest ELBO was selected and the Pearson correlation matrix between the factors is shown. **C:** Factor values are shown for all samples as points and coloured based on sex (upper panel) or BMI (lower panel). P-values from linear mixed model.

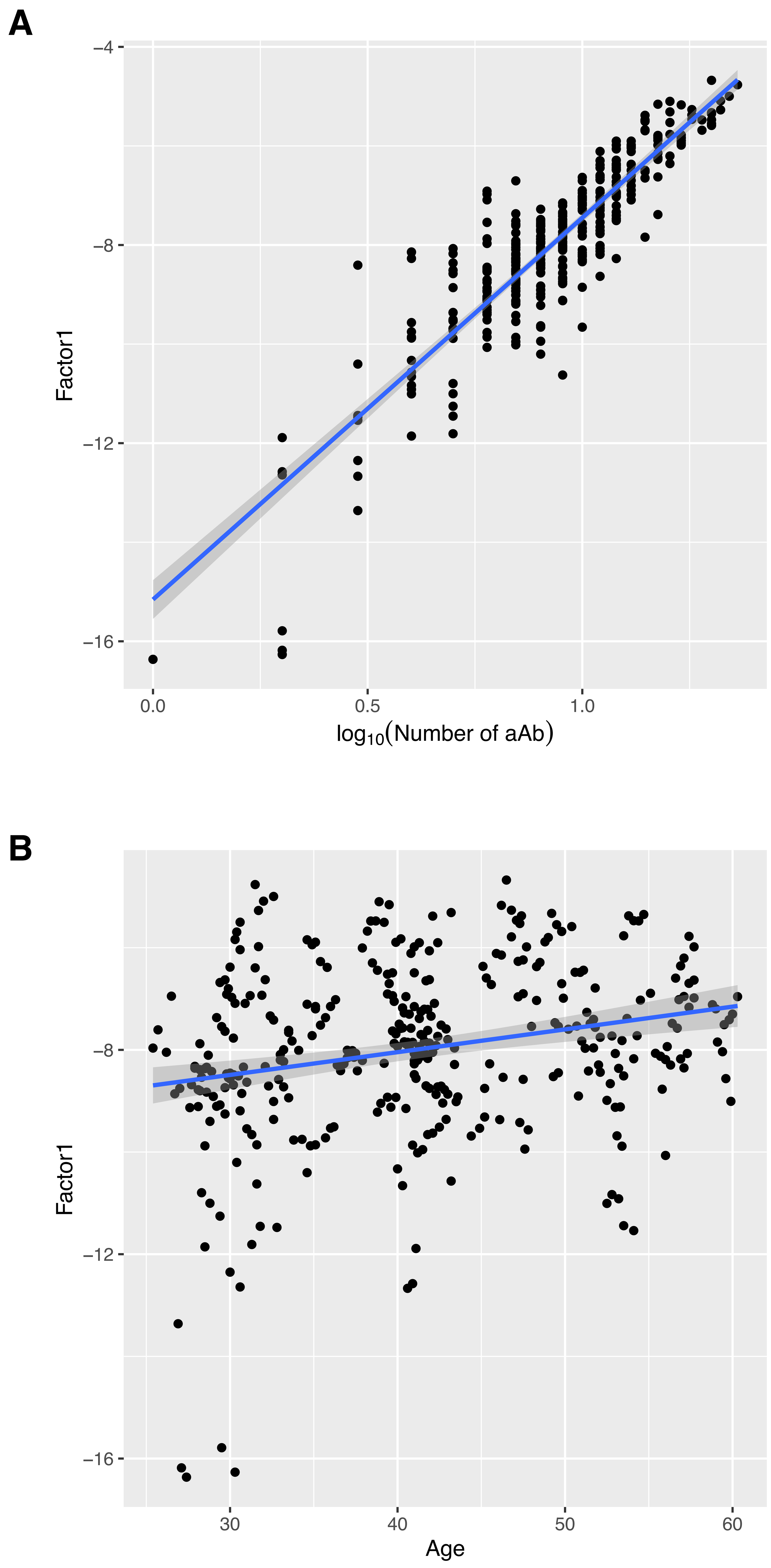


**Supplementary figure 8**: **A.** Factor 1 is associated with the number of detected autoantibodies. **B.** A scatterplot of the individual's age vs. Factor 1 values. Regression lines are shown with 95% confidence intervals.

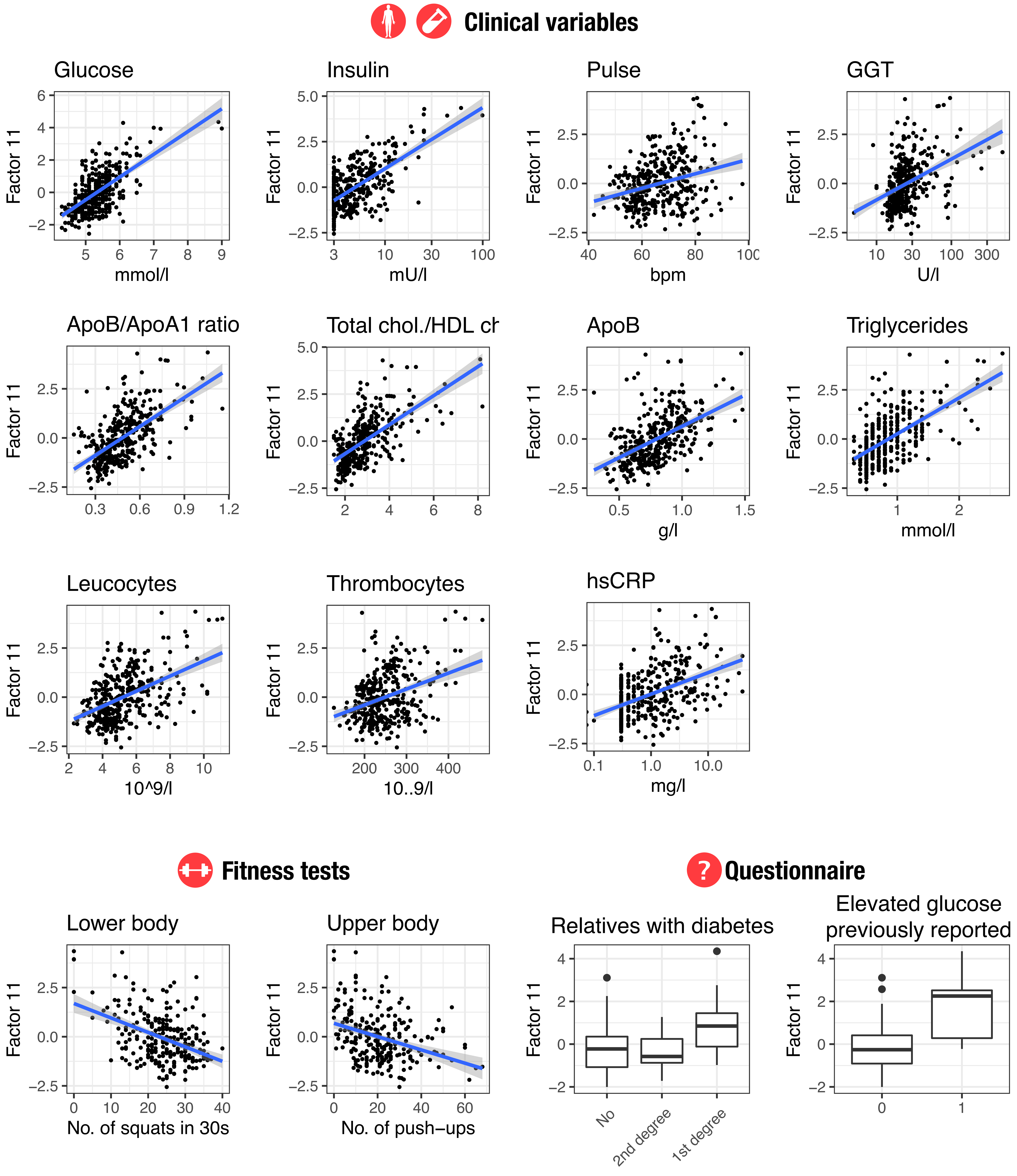


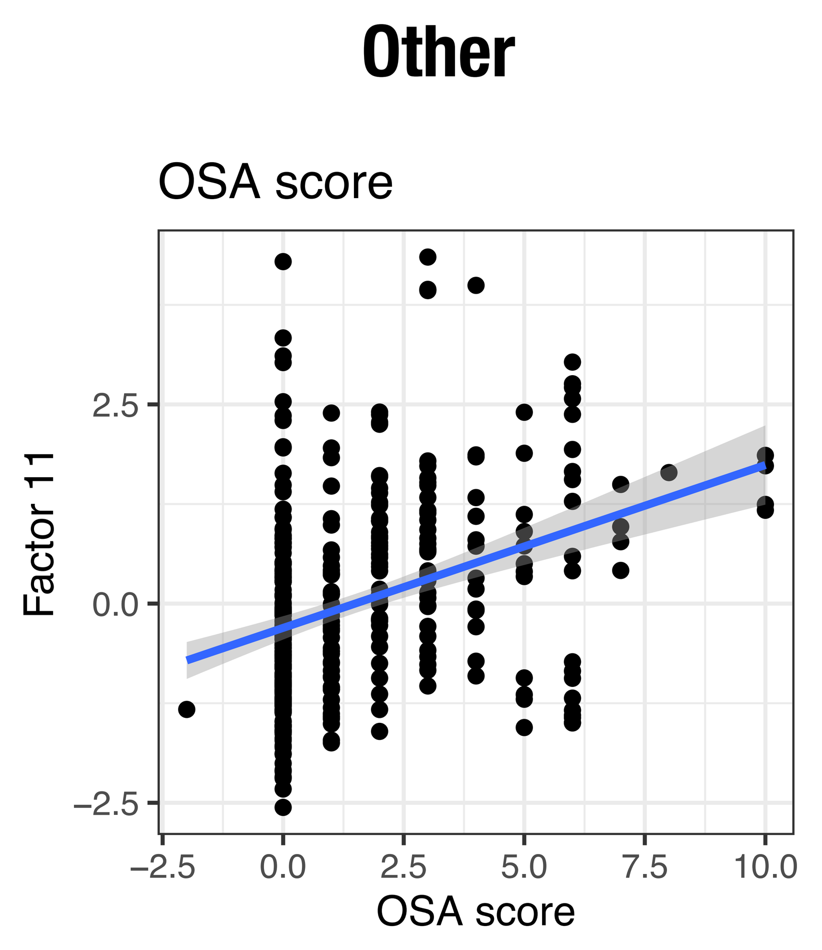


**Supplementary figure 9:** Additional plots showing the association of Factor11 with phenotypic variables. Regression lines are added when appropriate and 95% confidence intervals are shown. OSA: Obstructive Sleep Apnea

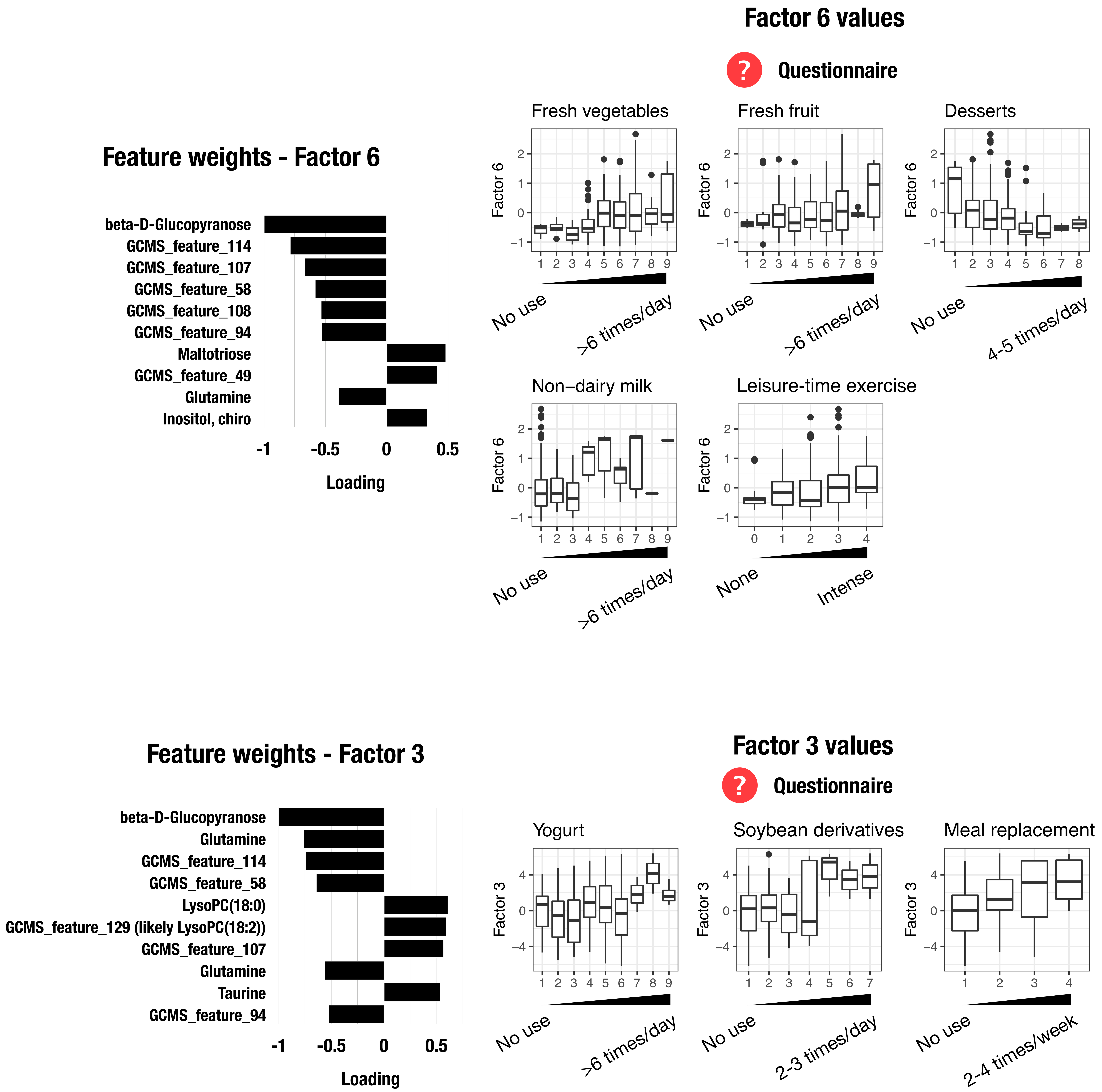


**Supplementary figure 10:** Factor 3 and Factor 6 are associated with diet. The top scaled feature weights are shown. The association between the factor values for each sample and the phenotypic variables is shown. A regression line with 95% confidence interval is shown when appropriate.

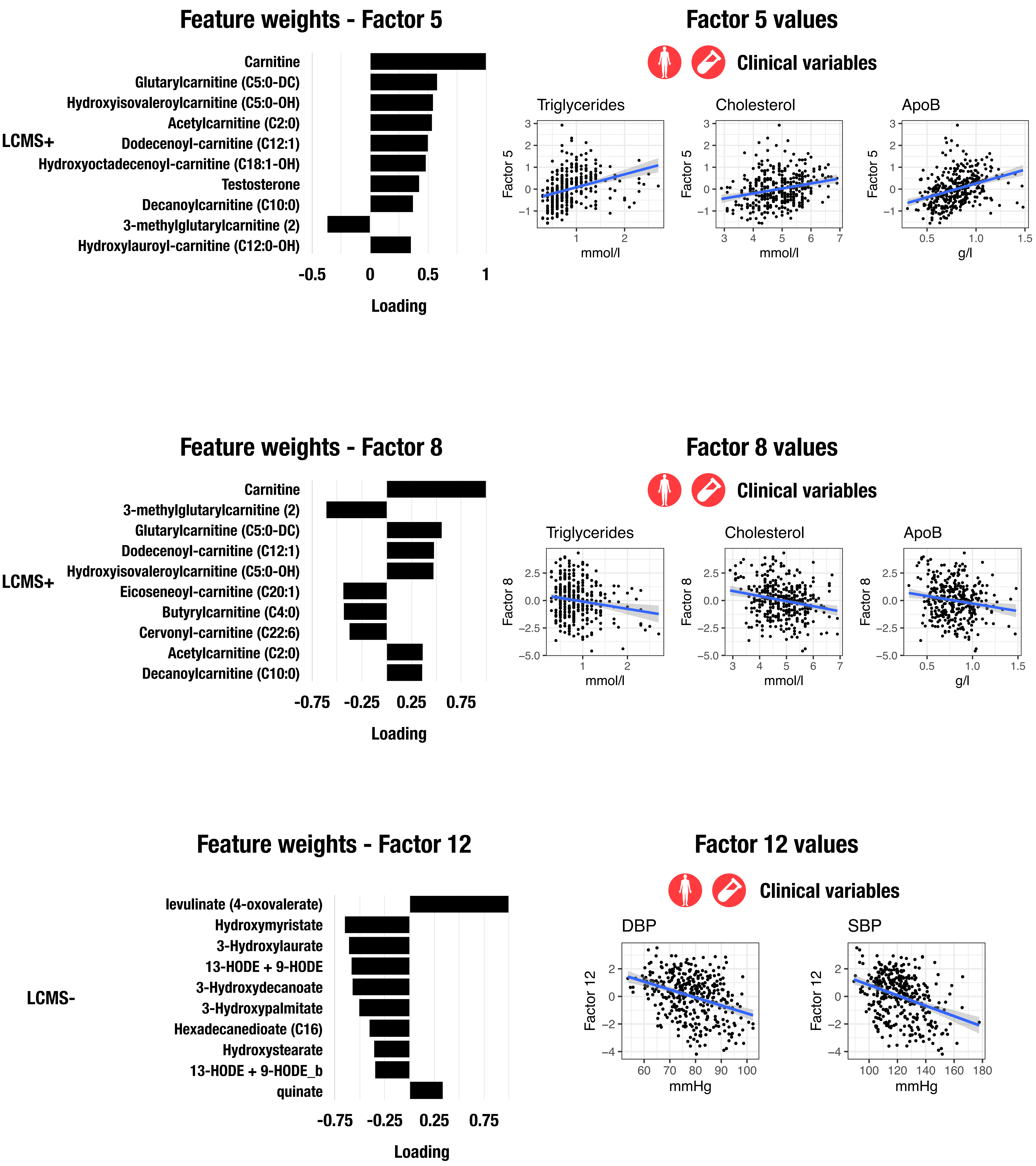


**Supplementary figure 11:** Factor 5, Factor 8 and Factor 12 are associated with fatty acids and acylcarnitines. The top scaled feature weights are shown. The association between the factor values for each sample and the phenotypic variables is shown. A regression line with 95% confidence interval is shown when appropriate.

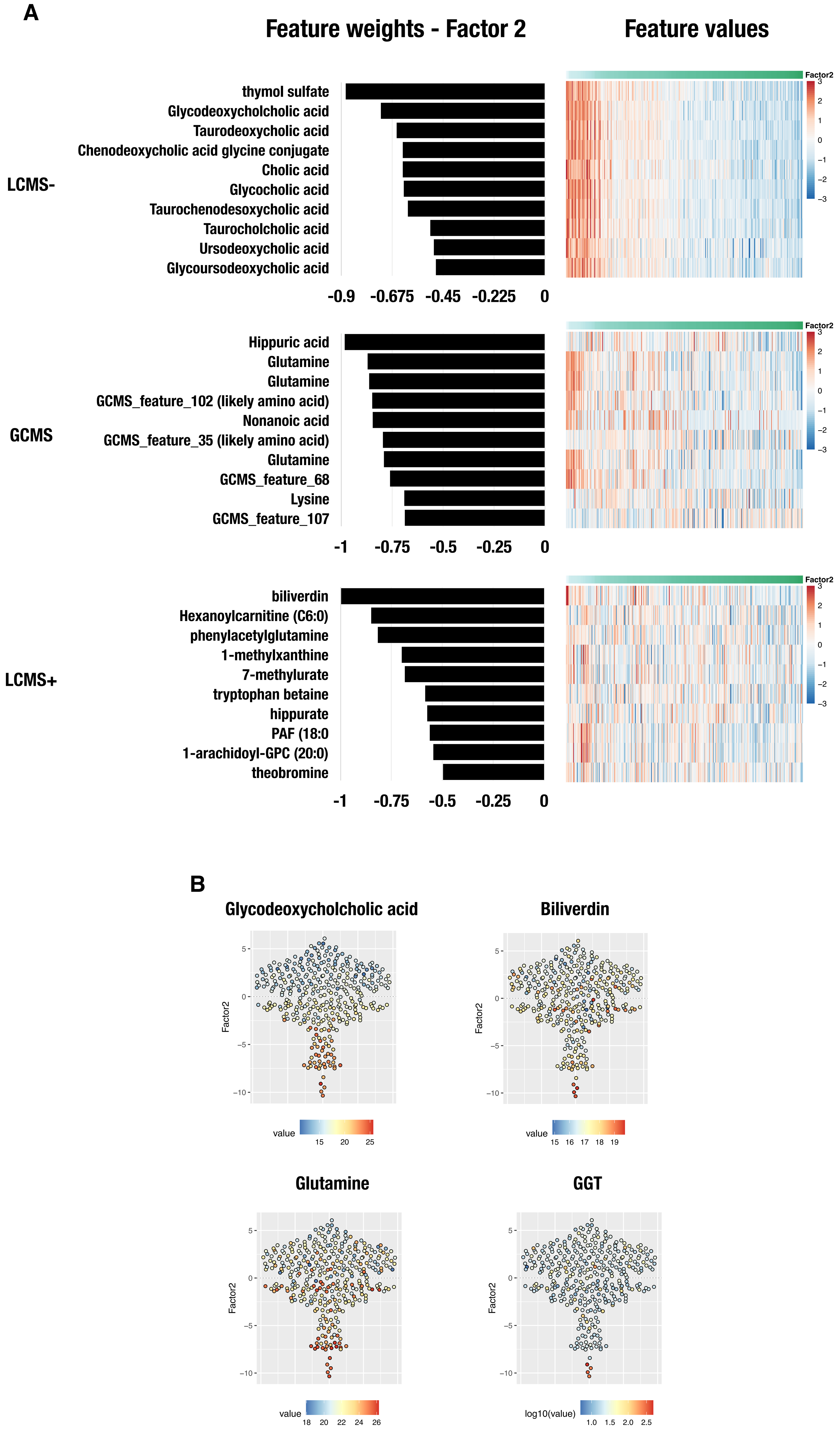
A
Feature weights - Factor 2
Feature values
LCMS-
thymol sulfate
Glycodeoxycholcholic acid
Taurodeoxycholic acid
Chenodeoxycholic acid glycine conjugate
Cholic acid
Glycocholic acid
Taurochenodesoxycholic acid
Taurocholcholic acid
Ursodeoxycholic acid
Glycoursodeoxycholic acid
-0.9 -0.675 -0.45 -0.225 0
GCMS
Hippuric acid
Glutamine
Glutamine
GCMS_feature_102 (likely amino acid)
Nonanoic acid
GCMS_feature_35 (likely amino acid)
Glutamine
GCMS_feature_68
Lysine
GCMS_feature_107
-1 -0.75 -0.5 -0.25 0
LCMS+
biliverdin
Hexanoylcarnitine (C6:0)
phenylacetylglutamine
1-methylxanthine
7-methylurate
tryptophan betaine
hippurate
PAF (18:0
1-arachidoyl-GPC (20:0)
theobromine
-1 -0.75 -0.5 -0.25 0
Factor2
B
Glycodeoxycholcholic acid
Biliverdin
Glutamine
GGT
Factor2
value
log10(value)

**Supplementary figure 12 (previous page): A.** Factor 2 is associated with hepatic function. The top negative scaled metabolite weights for Factor2 are shown, as well as their values (row z-score) as heatmaps. **B.** The Factor 2 values are shown for each sample and colored according to the value of the indicated feature.

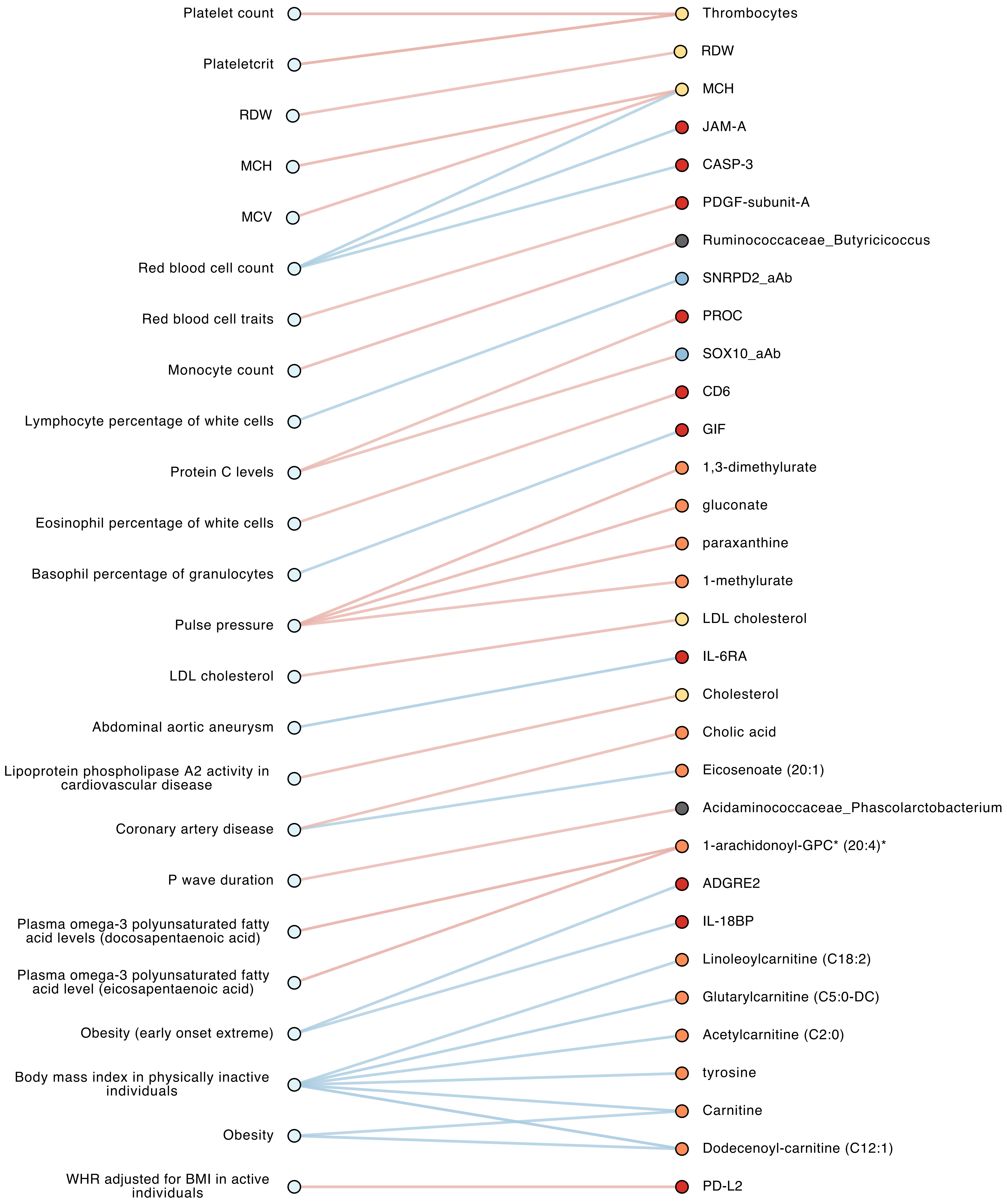


**Supplementary figure 13:** Association with genetic scores in the Between-Individual Network. All the edges between a GWAS summary score and another features were selected and shown here. Colors of the edges and nodes are the same as described in main Figure 5.

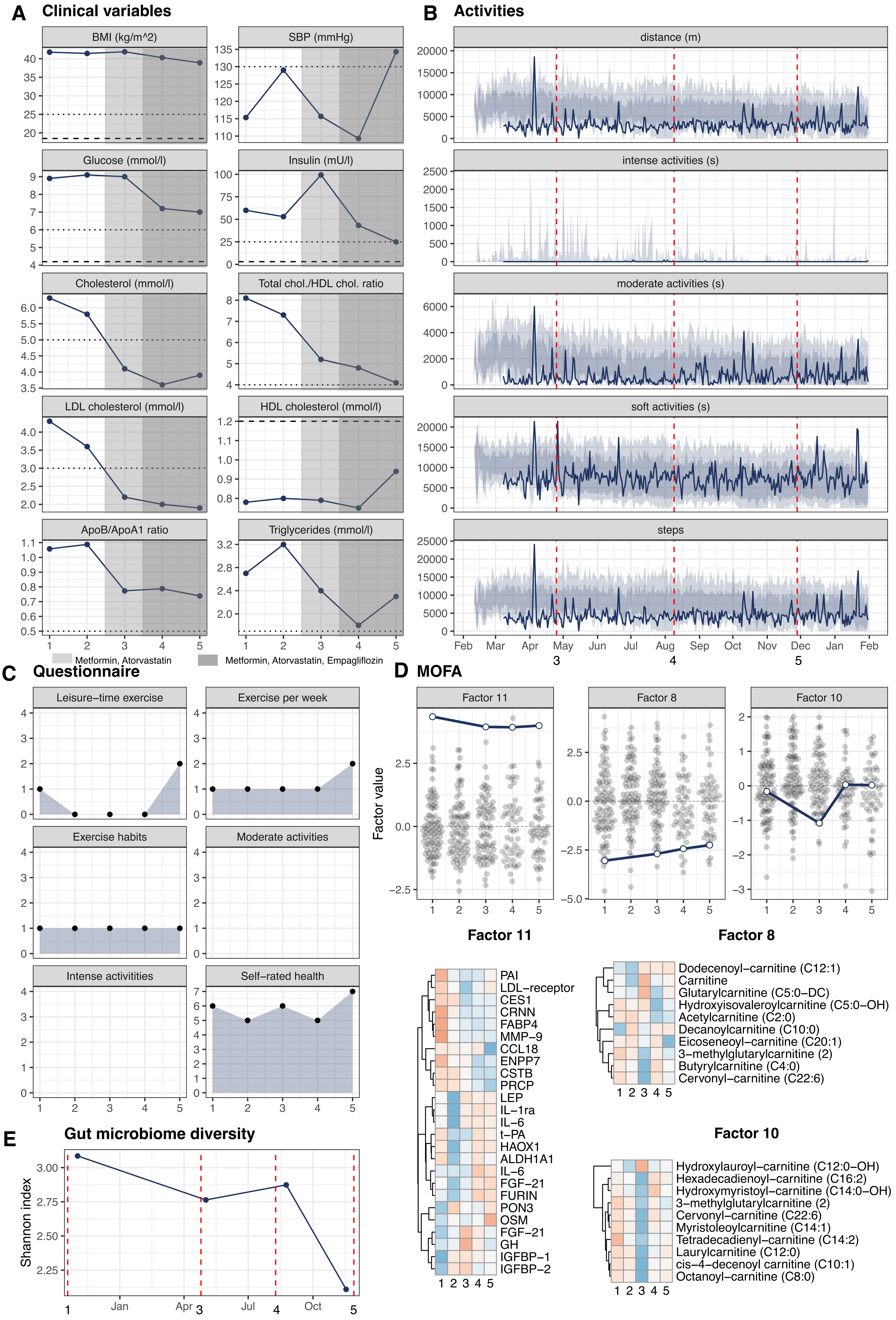

A Clinical variables
BMI (kg/m^2)
SBP (mmHg)
Glucose (mmol/l)
Insulin (mU/l)
Cholesterol (mmol/l)
Total chol./HDL chol. ratio
LDL cholesterol (mmol/l)
HDL cholesterol (mmol/l)
ApoB/ApoA1 ratio
Triglycerides (mmol/l)
Metformin, Atorvastatin
Metformin, Atorvastatin, Empagliflozin
B Activities
distance (m)
intense activities (s)
moderate activities (s)
soft activities (s)
steps
Feb Mar Apr May Jun Jul Aug Sep Oct Nov Dec Jan Feb
C Questionnaire
Leisure−time exercise
Exercise per week
Exercise habits
Moderate activities
Intense activitities
Self−rated health
D MOFA
Factor 11
Factor 8
Factor 10
Factor value
PAI
LDL−receptor
CES1
CRNN
FABP4
MMP−9
CCL18
ENPP7
CSTB
PRCP
LEP
IL−1ra
IL−6
t−PA
HAOX1
ALDH1A1
IL−6
FGF−21
FURIN
PON3
OSM
FGF−21
GH
IGFBP−1
IGFBP−2
Dodecenoyl−carnitine (C12:1)
Carnitine
Glutarylcarnitine (C5:0−DC)
Hydroxyisovaleroylcarnitine (C5:0−OH)
Acetylcarnitine (C2:0)
Decanoylcarnitine (C10:0)
Eicoseneoyl−carnitine (C20:1)
3−methylglutarylcarnitine (2)
Butyrylcarnitine (C4:0)
Cervonyl−carnitine (C22:6)
Hydroxylauroyl−carnitine (C12:0−OH)
Hexadecadienoyl−carnitine (C16:2)
Hydroxymyristoyl−carnitine (C14:0−OH)
3−methylglutarylcarnitine (2)
Cervonyl−carnitine (C22:6)
Myristoleoylcarnitine (C14:1)
Tetradecadienyl−carnitine (C14:2)
Laurylcarnitine (C12:0)
cis−4−decenoyl carnitine (C10:1)
Octanoyl−carnitine (C8:0)
E Gut microbiome diversity
Shannon index
Jan Apr Jul Oct

**Supplementary figure 14 (previous page):** A case study: one individual that was diagnosed with type 2 diabetes. **A.** Clinical parameters showing longitudinal changes. Reference values are shown (dashed line: lower normal value, dotted line: upper normal value). Grey-shaded areas mark periods of therapeutical intervention, consisting in Metformin and Atorvastatin before visit 2 and Metformin, Atorvastatin and Empagliflozin from visit 3 onward. **B.** Daily summary of the activities recorder by the smart watch included: distance, time spent in intense, moderate or soft activities and number of steps. The shaded area correspond to the 10, 25, 75 and 90 percentiles in the whole population. The study visits are marked with vertical dashed lines **C.** Selected lifestyle questions. The numbers correspond to orderer categories, where a higher number corresponds to higher frequency or more favorable outcome. **D.** Longitudinal values for selected MOFA+ factors (blue line) plotted together with the factor values for the remaining samples at each study visit (grey dots). The top features for the corresponding factors are shown as heatmaps (row z-score, blue:low, red:high). **E.** Gut microbiome diversity (Shannon index). The vertical dashed line mark the date for the study visit, which could be different than the fecal sample collection date (dots).

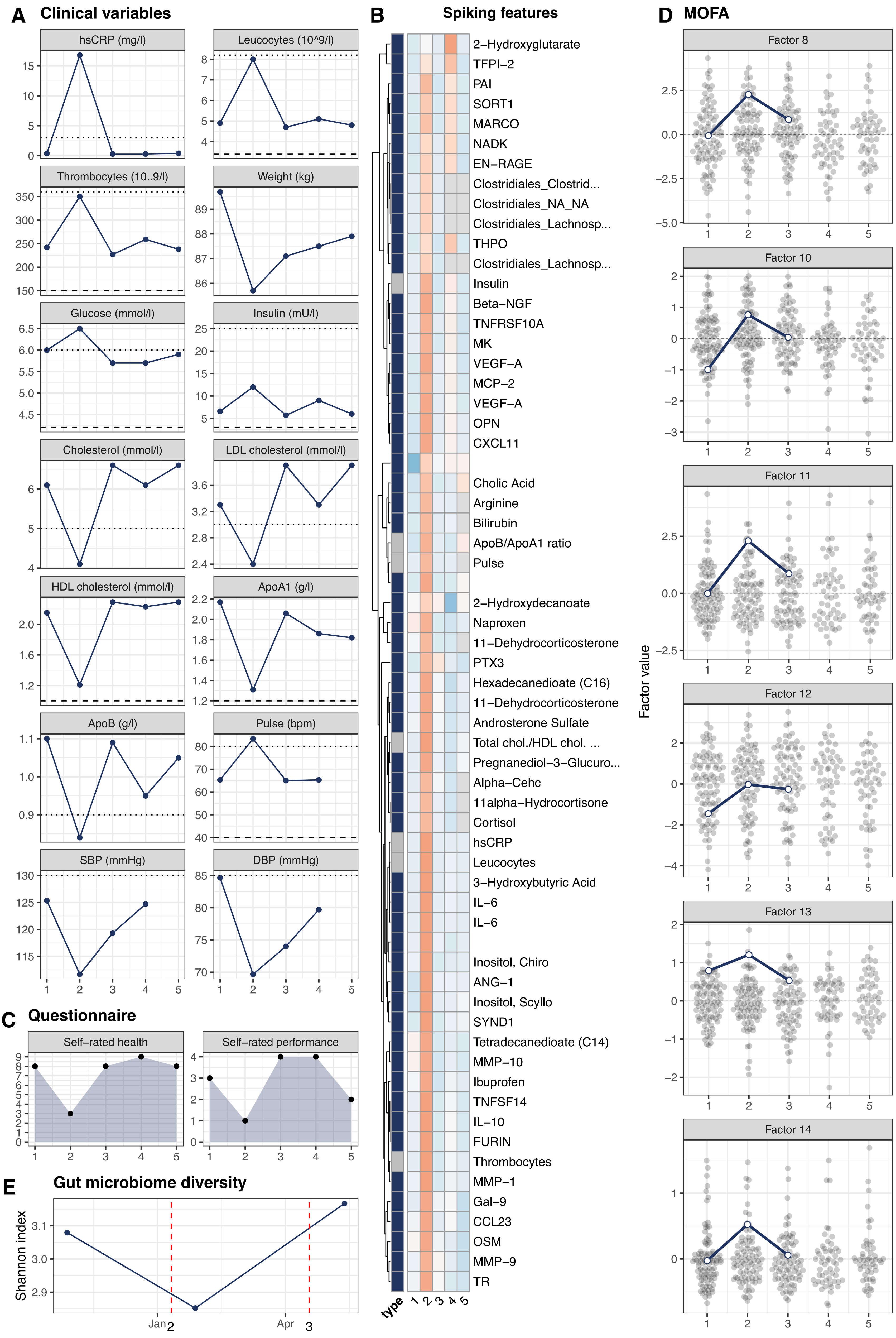
A Clinical variables
hsCRP (mg/l)
Leucocytes (10^9/l)
Thrombocytes (10..9/l)
Weight (kg)
Glucose (mmol/l)
Insulin (mU/l)
Cholesterol (mmol/l)
LDL cholesterol (mmol/l)
HDL cholesterol (mmol/l)
ApoA1 (g/l)
ApoB (g/l)
Pulse (bpm)
SBP (mmHg)
DBP (mmHg)
B Spiking features
2–Hydroxyglutarate
TFPI–2
PAI
SORT1
MARCO
NADK
EN–RAGE
Clostridiales_Clostrid...
Clostridiales_NA_NA
Clostridiales_Lachnosp...
THPO
Clostridiales_Lachnosp...
Insulin
Beta–NGF
TNFRSF10A
MK
VEGF–A
MCP–2
VEGF–A
OPN
CXCL11
Cholic Acid
Arginine
Bilirubin
ApoB/ApoA1 ratio
Pulse
2–Hydroxydecanoate
Naproxen
11–Dehydrocorticosterone
PTX3
Hexadecanedioate (C16)
11–Dehydrocorticosterone
Androsterone Sulfate
Total chol./HDL chol. ...
Pregnanediol–3–Glucuro...
Alpha–Cehc
11alpha–Hydrocortisone
Cortisol
hsCRP
Leucocytes
3–Hydroxybutyric Acid
IL–6
IL–6
Inositol, Chiro
ANG–1
Inositol, Scyllo
SYND1
Tetradecanedioate (C14)
MMP–10
Ibuprofen
TNFSF14
IL–10
FURIN
Thrombocytes
MMP–1
Gal–9
CCL23
OSM
MMP–9
TR
type
1 2 3 4 5
C Questionnaire
Self–rated health
Self–rated performance
D MOFA
Factor 8
Factor 10
Factor 11
Factor 12
Factor 13
Factor 14
Factor value
E Gut microbiome diversity
Shannon index
Jan
Apr
2
3

**Supplementary figure 15 (previous page).** A participant reported an unspecified respiratory tract infection at the second visit and presented with a ill-health phenotype, corresponding to a broad alteration of the molecular space suggested by changes in several MOFA+ factors at the second visit, including a drop in the Factors linked to acylcarnitines and lipid metabolism (F8, F10), a change in the circulating lipid pool (F12) and possible hormonal changes (F14). **A.** Clinical parameters showing altered values at baseline and longitudinal changes. Reference values are shown (dashed line: lower normal value, dotted line: upper normal value). **B.** The features with positive local maxima values (i.e. a peak) at the second visit are shown as a heatmap (row z-score). An annotation column indicates the feature type (blue: omics, grey: clinical variables). Local peaks were considered only if the peak height was larger than the median value plus 2.5 times the median absolute deviation value. The features revealed a response to infection/inflammation signature (cytokines, chemokines, growth factors and proteases among others), increased circulation of certain FFAs, and steroid hormones changes. An elevation of blood glucose and a small change in insulin levels were also detected. The levels of anti-inflammatory drug naproxen, ibuprofen also reached a local maximum at visit two **C.** Selected lifestyle questions. The numbers correspond to orderer categories, where a higher number corresponds to a more favorable outcome. **D.** Longitudinal values for selected MOFA+ factors (blue line) plotted together with the factor values for the remaining samples at each study visit (grey dots). The broad alteration of the molecular space at the second visit was suggested by changes in several MOFA+ factors (F8, F10, F12 and F14 - acylcarnitines, lipid metabolism, fatty acids and hormonal changes). **E.** Gut microbiome diversity (Shannon index). The vertical dashed line mark the date for the study visit, which could be different than the fecal sample collection date (dots). A decrease in gut microbiome diversity was observed, accompanied by an increase in the abundance of taxa of the Clostridia class.
